# AAV-mediated SIL1 gene therapy prevents cerebellar degeneration and ataxia in a mouse model of Marinesco-Sjögren syndrome

**DOI:** 10.64898/2026.09.14.751587

**Authors:** Chiara Pasini, Giada Lavigna, Anna Grasso, Elena Restelli, Alessandro Corbelli, Monica Salio, Lorena Zentilin, Fabio Fiordaliso, Roberto Chiesa

**Affiliations:** Laboratory of Prion Neurobiology, Department of Neuroscience, Istituto di Ricerche Farmacologiche Mario Negri IRCCS; Milan, 20156, Italy; Unit of Cellular Pathology, Department of Biochemistry and Molecular Pharmacology, Istituto di Ricerche Farmacologiche Mario Negri IRCCS; Milan, 20156, Italy; AAV Vector Unit, International Centre for Genetic Engineering and Biotechnology (ICGEB); Trieste, 34149, Italy

**Keywords:** Marinesco-Sjögren syndrome, SIL1, adeno-associated virus (AAV) gene therapy, cerebellar ataxia, Purkinje cells, endoplasmic reticulum stress

## Abstract

Marinesco-Sjögren syndrome is a rare early-onset multisystem disorder characterized primarily by cerebellar ataxia and myopathy and caused by loss-of-function mutations in *SIL1*. No disease-modifying therapy is available. We investigated whether adeno-associated virus (AAV)-mediated gene therapy could prevent neurological and muscular disease in the woozy mouse model of SIL1 deficiency. Neonatal mice received intracerebroventricular injections of AAV-PHP.eB vectors expressing SIL1 under the control of either a ubiquitous or a Purkinje cell-specific promoter. Vehicle-treated woozy mice developed progressive motor impairment accompanied by extensive Purkinje cell degeneration, thinning of the cerebellar molecular layer, marked astrogliosis and activation of endoplasmic reticulum stress pathways. In contrast, AAV-SIL1 treatment prevented the onset of ataxia and substantially preserved cerebellar architecture, including Purkinje cells and molecular-layer thickness, while reducing astrogliosis and endoplasmic reticulum stress. Selective restoration of SIL1 expression in Purkinje cells was sufficient to rescue motor performance, supporting a major cell-autonomous contribution of Purkinje cell dysfunction to cerebellar disease. Therapeutic benefit was maintained throughout the 26-week observation period, the longest time point examined, with treated mice remaining behaviorally indistinguishable from heterozygous controls. AAV-SIL1 treatment also improved muscle function and markedly attenuated skeletal muscle pathology, with transgenic SIL1 expression in muscle reaching levels comparable to those in heterozygous controls. These findings provide proof-of-principle that AAV-mediated SIL1 gene therapy can prevent the neurological and muscular manifestations of Marinesco-Sjögren syndrome and identify Purkinje cells as a critical cellular target for preventing cerebellar dysfunction.

## INTRODUCTION

Marinesco-Sjögren syndrome (MSS) is a rare autosomal recessive disorder with onset in early infancy and no curative treatment. Its cardinal features are congenital cataracts, cerebellar atrophy and myopathy, with additional manifestations including intellectual disability, intention tremor, dysarthria, skeletal abnormalities, short stature and hypogonadism ^1,2^. Initial symptoms typically appear within the first months of life, as affected infants fail to achieve early motor milestones. Motor function typically deteriorates during childhood but, after a period of progression lasting several years, the disease generally stabilizes at an unpredictable age and degree of severity, and life expectancy is normal ^1^. Therefore, interventions that prevent, delay, or attenuate disease progression during the early phase of the disorder could have a profound and lasting impact on motor function and quality of life.

Approximately 60% of MSS cases are caused by homozygous or compound heterozygous loss-of-function mutations in *SIL1*, resulting in reduced or absent SIL1 protein function ^3,4^. A spontaneous *Sil1* mutation underlies the phenotype of woozy (wz) mice, which recapitulate key features of MSS, including cerebellar ataxia with Purkinje cell (PC) degeneration and skeletal muscle pathology ^5,6^.

SIL1 encodes an endoplasmic reticulum (ER) nucleotide exchange factor for the molecular chaperone BiP. By facilitating ADP-ATP exchange, SIL1 promotes the release and refolding of BiP-bound client proteins. SIL1 deficiency disrupts this process, leading to accumulation of unfolded proteins, ER stress and activation of the unfolded protein response (UPR) ^2^.

Among the three major UPR pathways, PERK phosphorylates eukaryotic translation initiation factor 2α (eIF2α), reducing global protein synthesis while promoting translation of ATF4. Sustained ATF4 activation induces CHOP and can ultimately trigger apoptosis ^7^. This pathway is particularly detrimental to neurons, which rely heavily on ongoing protein synthesis for synaptic maintenance and survival ^8^.

We previously showed that PERK/eIF2α signalling precedes PC degeneration and ataxia in woozy mice and is activated in skeletal muscle at the onset of myopathy ^9,10^. Early treatment with the PERK inhibitor GSK2606414 delayed neurodegeneration and motor deficits and ameliorated muscle pathology, but was associated with pancreatic toxicity and did not fully rescue the phenotype ^9,11^. Other approaches targeting PERK signalling downstream of eIF2α, including dibenzoylmethane and trazodone, as well as the chemical chaperone tauroursodeoxycholic acid, also failed to prevent motor deficits or PC loss in woozy mice ^12^.

Because MSS is primarily caused by loss of SIL1 function, restoring the deficient protein represents a more direct therapeutic strategy than targeting downstream consequences of ER dysfunction. Recombinant adeno-associated virus (AAV) vectors are well suited for this purpose because they can provide sustained transgene expression in post-mitotic tissues, including the CNS and skeletal muscle, and have an established clinical safety profile ^13^. Advances in capsid engineering have further improved the efficiency of CNS gene delivery.

Here, we evaluated whether neonatal intracerebroventricular (icv) administration of AAV vectors expressing SIL1 could prevent cerebellar and skeletal muscle pathology and provide sustained functional benefit in the woozy mouse model of MSS. We further tested whether selective restoration of SIL1 expression in PCs was sufficient to rescue cerebellar dysfunction, thereby investigating the contribution of this neuronal population to MSS pathogenesis.

## RESULTS

### AAV vectors drive efficient and cell-selective SIL1 expression with correct endoplasmic reticulum localization

We generated AAV vectors encoding mouse (mo) or human (hu) SIL1 under the ubiquitous CAG promoter by replacing GFP in pAAV-CAG-GFP with the corresponding SIL1 cDNAs. To test whether selective restoration of SIL1 in Purkinje cells is sufficient to rescue cerebellar dysfunction, we also generated vectors expressing moSIL1 or GFP under the PC-specific L7-6 promoter ^14^.

AAV9 and AAV-PHP.eB (PHP.eB) capsids were first tested in woozy mouse embryonic fibroblasts (MEFs), which lack endogenous SIL1. Both AAV9-CAG-moSIL1 and PHP.eB-CAG-moSIL1 induced SIL1 expression, with PHP.eB showing higher efficiency, whereas AAV9-L7-6-moSIL1 did not, consistent with the PC specificity of the L7-6 promoter (Fig. S1A). Transgenic SIL1 co-localized with the ER marker PDI, but not the Golgi marker GM130 (Fig. S1B,C), indicating correct ER localization. The AAV-encoded SIL1 protein was sensitive to deglycosylation with both PNGase F and Endo H, the latter indicating the presence of immature high-mannose N-glycans characteristic of proteins retained within the ER (Fig. S2).

We next confirmed L7-6 promoter specificity in cultured organotypic cerebellar slices (COCS). AAV9-CAG-GFP produced widespread GFP expression, whereas AAV9-L7-6-GFP expression was restricted to calbindin-positive PCs (Fig. S3). Consistent with selective PC expression, PHP.eB-L7-6-SIL1 produced markedly lower SIL1 levels than PHP.eB-CAG-SIL1 in woozy COCS (Fig. S4).

### Intracerebroventricular delivery of AAVs encoding SIL1, either ubiquitously or selectively in Purkinje cells, reduces CHOP-positive PCs, mitigates early degeneration, and prevents ataxia

Woozy mice develop detectable ataxia by 6–7 weeks on the beam-walking test and by 9–10 weeks on the accelerating rotarod. Motor performance progressively declines until 16 weeks, when it stabilizes, while PC degeneration begins at 7–8 weeks and is nearly complete by 16 weeks ^9^. We therefore first tested whether neonatal AAV administration could prevent early motor dysfunction and PC degeneration (Fig. S5A).

Intracerebroventricular delivery produced widespread CNS expression when GFP was driven by the CAG promoter and selective PC expression with the L7-6 promoter (Fig. S6A). Consistent with previous reports ^15–17^, GFP was predominantly neuronal (Fig. S7) and, in the cerebellum, largely restricted to PCs even with the CAG promoter (Fig. S6). Because PHP.eB provided higher transduction efficiency than AAV9 at equivalent viral titres (Fig. S6), it was used for subsequent experiments.

Heterozygous (CT) and woozy littermates received 10^11^ vg/mouse of PHP.eB-CAG-moSIL1, PHP.eB-L7-6-moSIL1, or PBS by icv injection at P0 and were assessed weekly by beam walking until 9 weeks (Fig. S5A). Additional animals treated with PBS or PHP.eB-CAG-moSIL1 were analysed at 6 weeks for UPR markers in laser-capture microdissected PCs, when UPR activation is maximal in untreated woozy mice ^9,12^.

Beam-walking performance did not differ among groups before 6 weeks. From 6 weeks onward, PBS-treated woozy mice showed significantly more missteps and longer crossing times than CT mice, whereas woozy mice treated with either CAG- or L7-6-SIL1 showed robust and comparable preservation of motor performance, similar to CT mice throughout the study (Fig. 1A,B).

**Figure 1.**
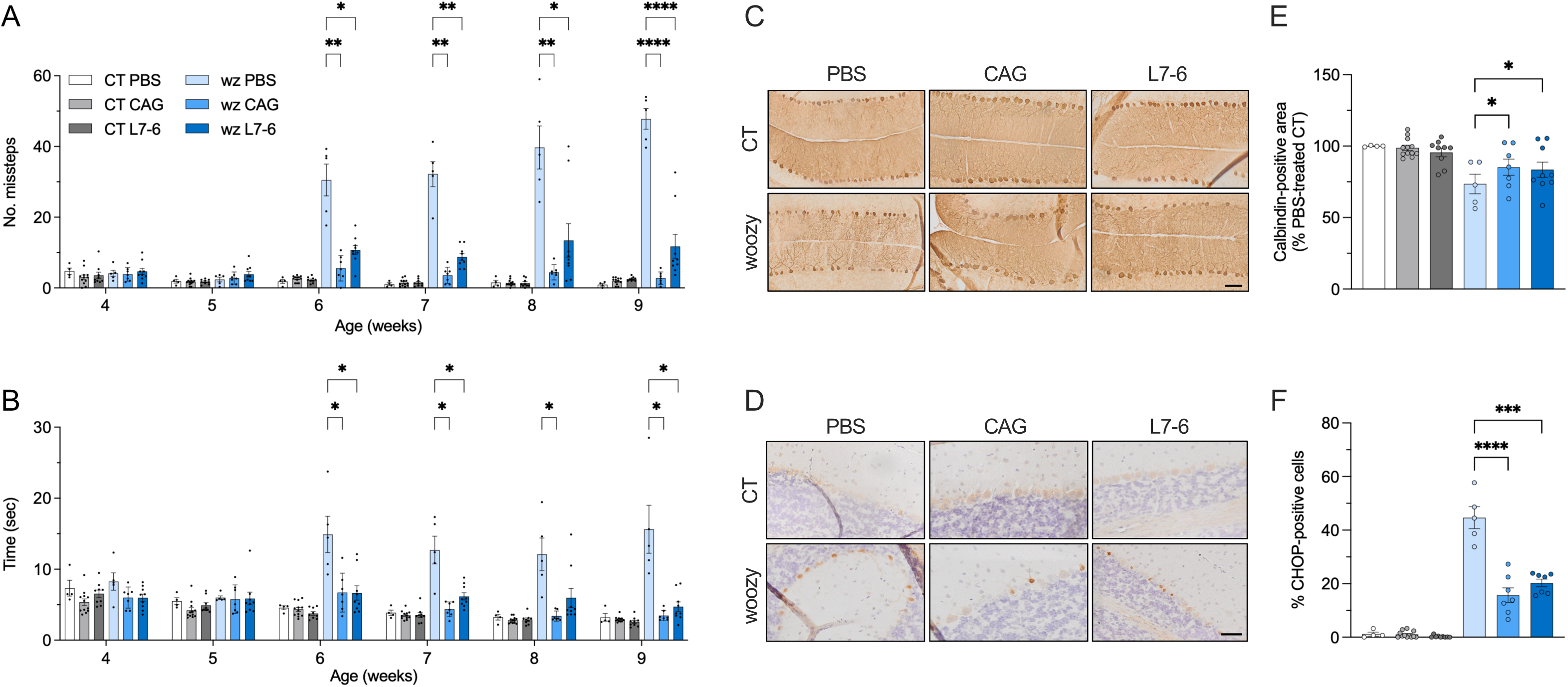
AAV-mediated SIL1 expression ameliorates motor performance and cerebellar pathology in woozy mice when expressed ubiquitously or selectively in PCs. (**A, B**) Groups of heterozygous control (CT) and woozy (wz) mice were injected icv with PBS, AAV-CAG-moSIL1 or AAV-L7-6-moSIL1. Animals were tested weekly for their ability to walk on a suspended beam. Data are presented as the mean ± SEM of the number of hindfoot missteps (A) and of the mean time to cross the beam (B). In (A), F(3.543, 31.89) = 17.66, *P* < 0.0001; in (B), F(4.926, 44.34) = 5.775, *P* = 0.0004 by two-way RM ANOVA; \*\**P* < 0.01, \*\**P* < 0.05, \*\*\*\**P* < 0.0001 by Dunnett’s post-hoc test. (**C, D**) Immunohistochemical analysis for calbindin (C) or CHOP (D) in cerebellar sections of CT and woozy mice at 9 weeks of age. Scale bar 200 μm. (**E, F**) Quantification of the calbindin-positive area in the cerebellar cortex (E), and of CHOP-positive PCs (F). Data are the mean ± SEM. In (E), F(2, 10) = 6.648, *P* = 0.0146; in (F), F(2, 9) = 33.35, *P* < 0.0001 by two-way ANOVA; \**P* < 0.05, \*\*\**P* < 0.001, \*\*\*\**P* < 0.0001 by Dunnett’s post-hoc test.

At 9 weeks, PBS-treated woozy mice showed a 27% reduction in calbindin-positive PC area compared with CT mice, whereas both CAG- and L7-6-SIL1 treatment significantly preserved PC area (Fig. 1C,E). Both treatments also reduced the proportion of CHOP-positive PCs (Fig. 1D,F). Consistent with these findings, ddPCR analysis of laser-capture microdissected PCs at 6 weeks showed reduced BiP, ATF4 and CHOP mRNA expression following CAG-SIL1 treatment (Fig. S8).

Western blotting confirmed robust transgenic SIL1 expression in the cerebrum and cerebellum following PHP.eB-CAG-moSIL1, whereas expression after PHP.eB-L7-6-moSIL1 was restricted to the cerebellum and substantially lower, consistent with PC-selective expression (Fig. S9). In CAG-moSIL1-treated PCs, transgenic SIL1 mRNA was approximately fourfold higher than in vehicle-treated CT mice, corresponding to an estimated twofold increase over endogenous SIL1 levels (Fig. S8).

We next tested whether human SIL1 produced a similar therapeutic effect and included PHP.eB-CAG-GFP as a control for vector administration. GFP-treated woozy mice developed progressive beam-walking deficits from 6–7 weeks, whereas mice treated with either moSIL1 or huSIL1 maintained performance comparable to CT mice (Fig. 2A,B). Both SIL1 vectors also increased PC preservation and reduced the proportion of CHOP-positive PCs at 9 weeks (Fig. 2C–F), confirming that the protective effect was attributable to SIL1 expression.

**Figure 2.**
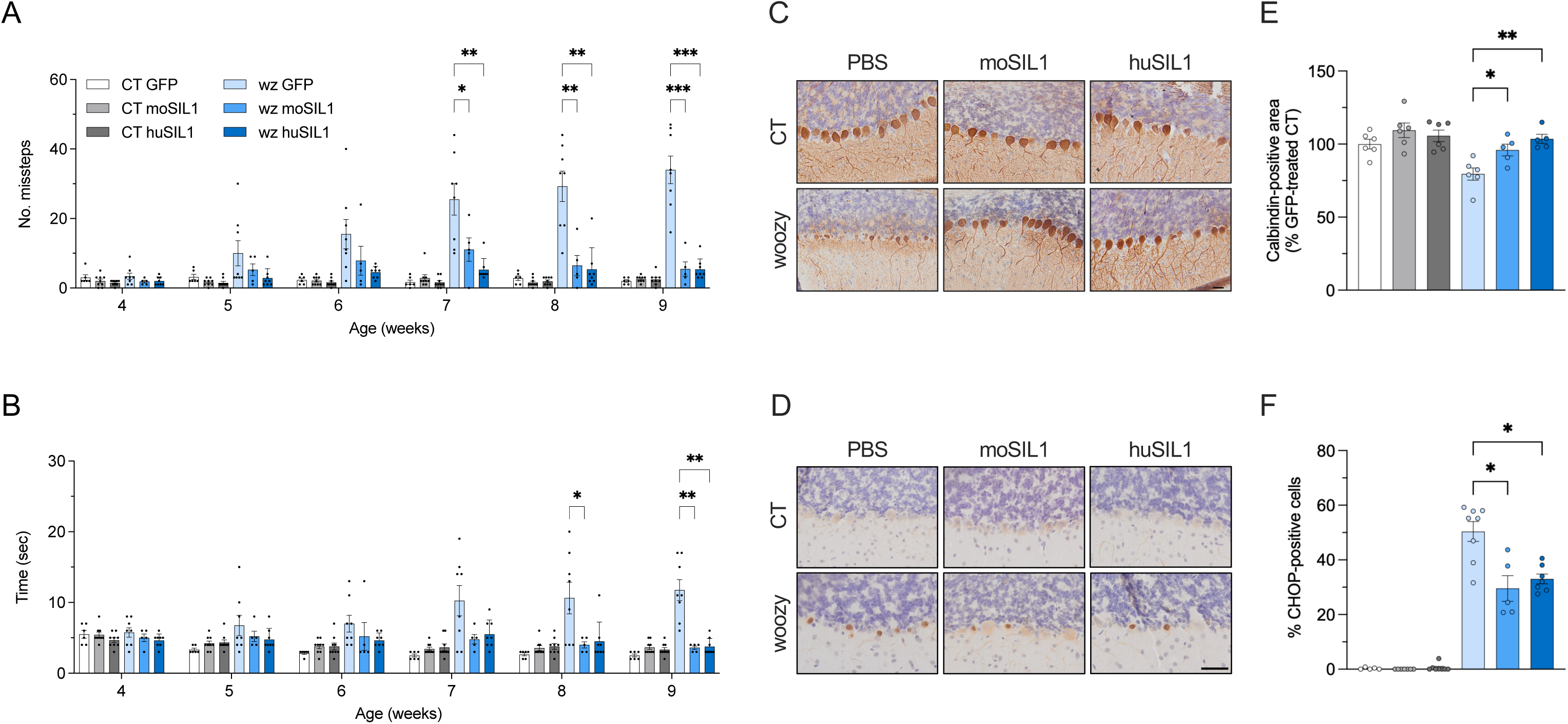
Neonatal intracerebroventricular injection of AAVs encoding either mouse or human SIL1 prevents development of motor deficits and PC degeneration in woozy mice. (**A, B**) Newborn woozy and heterozygous (CT) littermates were injected icv with PHP.eB-CAG-GFP, PHP.eB-CAG-moSIL1 or PHP.eB-CAG-huSIL1 (10^11^ vg/mouse), and tested weekly for their ability to walk on a suspended beam starting from four weeks of age. Data are the mean ± SEM of the number of hindfoot missteps (A) and the time required to cross the beam (B). In (A), F(5.605, 50.44) = 8.614, *P* < 0.0001; in (B), F(5.356, 48.21) = 3.966, *P* = 0.0036 by two-way RM ANOVA; \**P* < 0.05, \*\**P* < 0.01, \*\*\**P* < 0.001 by Dunnett’s post-hoc test. (**C, D**) Immunohistochemical analysis for calbindin (C) or CHOP (D) in cerebellar sections of CT and woozy mice at 9 weeks of age. Scale bar 200 μm. (**E, F**) Quantification of the calbindin-positive area in the cerebellar cortex (E), and of CHOP-positive PCs (F). Data are the mean ± SEM. In (E), F(2, 8) = 12.50, *P* = 0.0035; in (F), F(2, 10) = 6.942, *P* = 0.0129 by two-way ANOVA; \**P* < 0.05, \*\**P* < 0.01 by Dunnet’s post-hoc test.

Cerebellar SIL1 levels in moSIL1-treated woozy mice were approximately 1.5-fold those of endogenous wild-type mice, whereas huSIL1 produced approximately sixfold overexpression (Fig. 3).

**Figure 3.**
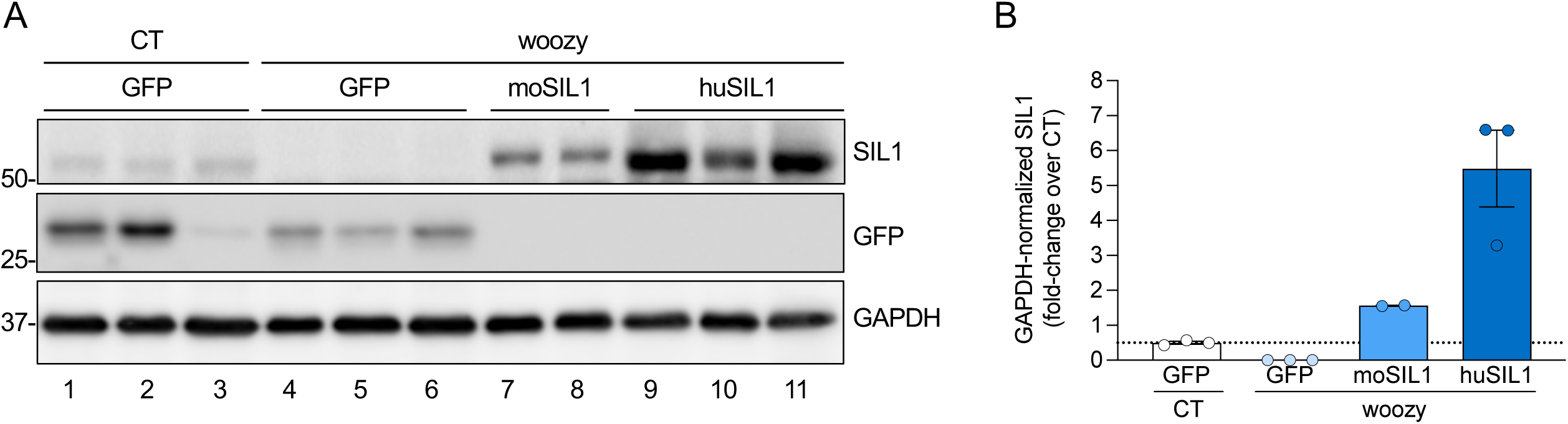
Intracerebroventricular injection of PHP.eB-CAG-huSIL1 and PHP.eB-CAG-moSIL1 induces robust transgene expression in the cerebellum. (**A**) Western blot analysis of SIL1, GFP and GAPDH in cerebellar homogenates of CT and woozy mice injected icv at P0 with PHP.eB-CAG-GFP, PHP.eB-CAG-huSIL1 or PHP.eB-CAG-moSIL1 at 9 weeks of age. (**B**) SIL1 levels were quantified by densitometry, normalized for the level of GAPDH, and plotted relative to the amount of endogenous SIL1 in PHP.eB-CAG-GFP-injected CT mice. Data are the mean ± SEM of 2-3 animals per group.

### AAVs encoding mouse or human SIL1 provide long-term protection against cerebellar ataxia and Purkinje cell degeneration

To determine whether the therapeutic effects of AAV treatment persisted long-term, we repeated the experiment, monitoring animals up to 26 weeks of age (Fig. S5B). Beam-walking performance during the first 9 weeks confirmed the protective effect of both moSIL1 and huSIL1 treatment (Fig. 4A,B; Movie S1).

**Figure 4.**
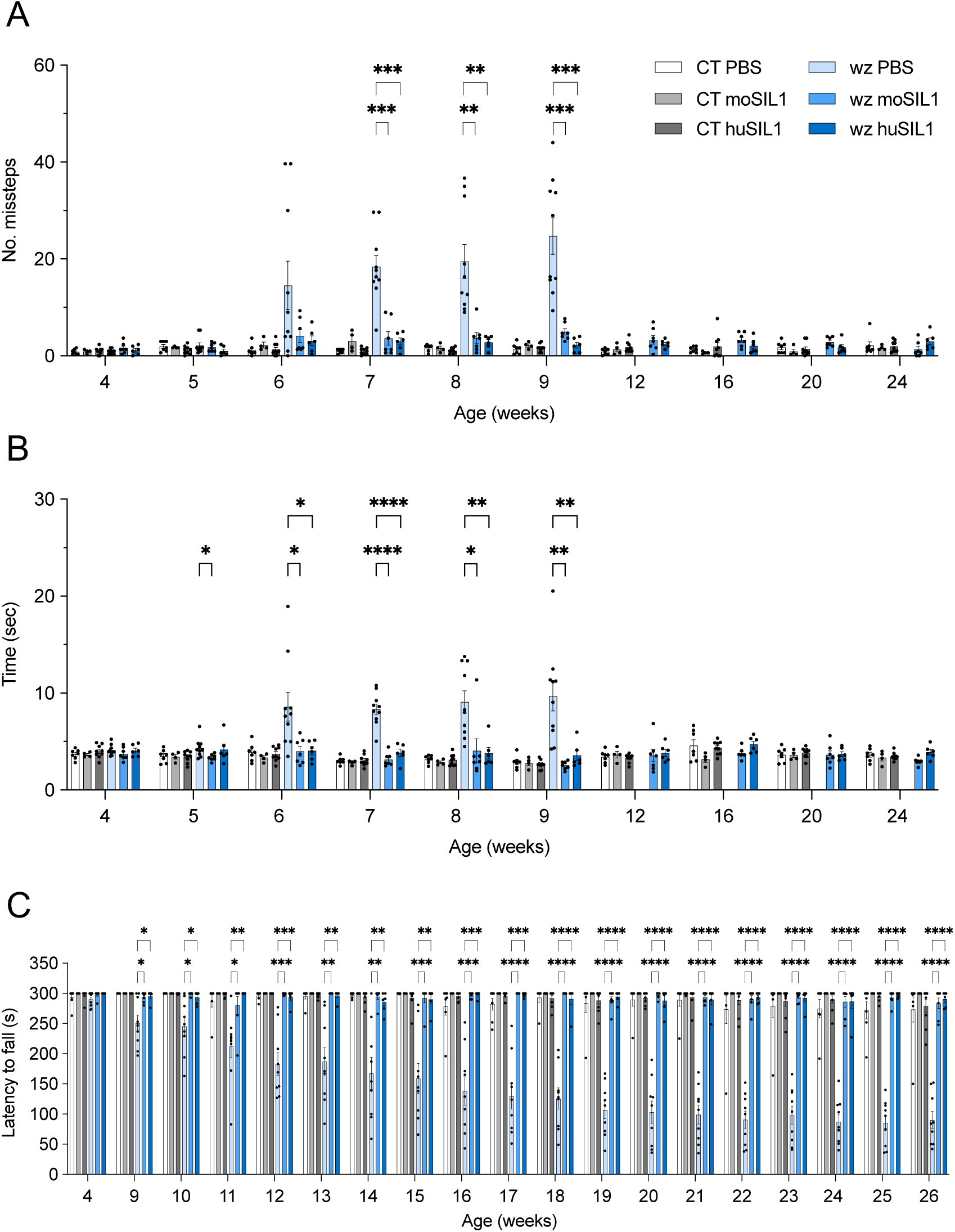
Intracerebroventricular injection of PHP.eB-CAG-moSIL1 and PHP.eB-CAG-huSIL1 in newborn woozy provides sustained protection from motor dysfunction. (**A**) Newborn woozy and heterozygous (CT) littermates were injected icv with PBS, PHP.eB-CAG-moSIL1 or PHP.eB-CAG-huSIL1, and tested for their ability to walk on a suspended beam starting from four weeks of age. Mice were tested weekly up to 9 weeks of age, thereafter PBS-treated woozy mice became unable to perform the test. Mice belonging to the other groups we retested at 12, 16, 20 and 24 weeks of age. Bars indicate the mean ± SEM of the number of hindfoot missteps (A) and the time required to cross the beam (B). In (A), F(4.420, 44.20) = 7.994, *P* < 0.0001; in (B) F (6.050, 60.50) = 5.038, *P* = 0.0003 by two-way RM ANOVA (1-9 weeks); \**P* < 0.05, \*\**P* < 0.01, \*\*\**P* < 0.001, \*\*\*\**P* < 0.0001 by Dunnett’s post-hoc test; (**C**) The same animals were tested at 4 weeks of age (baseline) and then weekly from 9 weeks onward on the accelerating rotarod. Bars indicate mean ± SEM latency to fall (s). F(3.169, 30.10) = 26.62, *P* < 0.0001; \**P* < 0.05, \*\**P* < 0.01, \*\*\**P* < 0.001, \*\*\*\**P* < 0.0001 by Dunnett’s post hoc test. Legend in (A) applicable to all panels.

Thereafter, vehicle-treated woozy mice became unable to perform the test, whereas AAV-treated woozy mice continued to walk on the beam without deterioration, with performance comparable to CT mice (Fig. 4A,B; Movie S2).

This long-term benefit was confirmed by the accelerated rotarod test, which is less demanding and can still be performed by woozy mice at advanced disease stages. Vehicle-treated woozy mice showed the expected progressive decline in latency to fall, whereas mice treated with either moSIL1 or huSIL1 maintained performance indistinguishable from CT mice throughout the study (Fig. 4C).

Progressive tremor is part of the neurological phenotype of woozy mice ^5,9^. Mild tremor is first apparent at 10 weeks of age when mice start walking on the suspended beam. As the mice age, the tremor becomes more severe and can also be seen when mice start walking on a table top ^9^. Woozy mice were scored for tremors at 20 and 26 weeks of age. At both time points, all nine mice in the vehicle-treated group had severe tremor, whereas none of the 13 AAV-treated mice showed any visible tremor, even when walking on the suspended beam (Movie S2).

At 26 weeks, calbindin immunohistochemistry showed complete PC loss and marked thinning of the cerebellar molecular layer in vehicle-treated woozy mice (Fig. 5). In contrast, AAV-treated woozy mice showed significant preservation of both PCs and molecular layer thickness. No detectable CHOP activation was observed in PCs of AAV-treated mice (Fig. S10), consistent with sustained suppression of ER stress.

**Figure 5.**
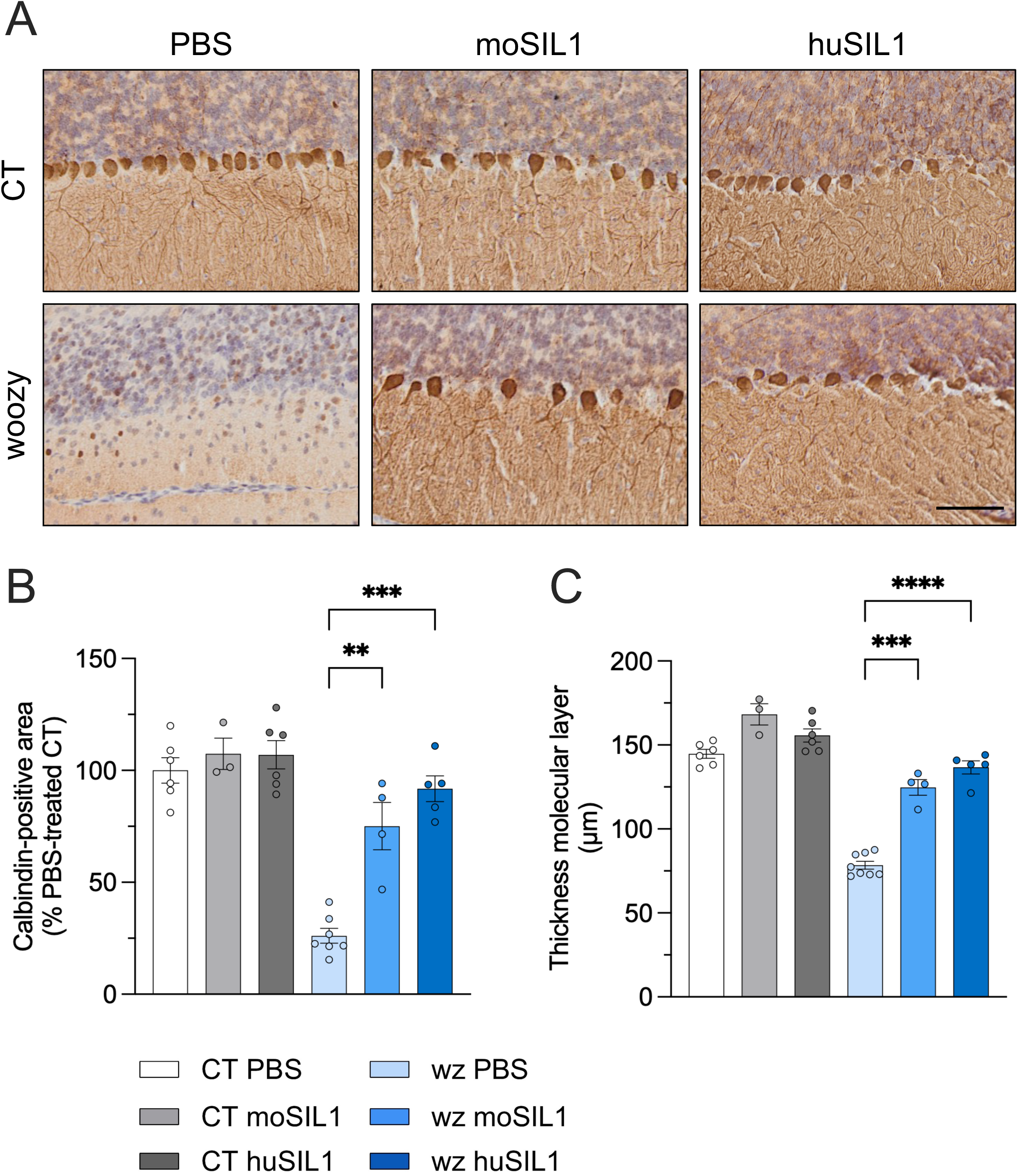
Intracerebroventricular injection of PHP.eB-CAG-moSIL1 and PHP.eB-CAG-huSIL1 rescues PCs degeneration in woozy mice. (**A**) Immunohistochemical analysis for calbindin in PBS, PHP.eB-CAG-moSIL1 or PHP.eB-CAG-huSIL1-treated CT or woozy mice at 26 weeks of age. Scale bar 100 μm. (**B**) Percentage of the calbindin positive area in the cerebellar cortex. Data are the mean ± SEM of 3 brain sections from 5-6 mice per experimental group. F(2, 7) = 20.98, *P* = 0.0011 by two-way ANOVA; \*\**P* < 0.01, \*\*\**P* < 0.001 by Dunnett’s post-hoc test. (**C**) Molecular layer thickness. Data are the mean ± SEM. F(2, 7) = 56.45, *P* < 0.0001 by two-way ANOVA; \*\*\**P* < 0.001, \*\*\*\**P* < 0.0001 by Dunnett’s post-hoc test.

Finally, GFAP immunostaining revealed marked astrogliosis throughout the cerebellar cortex of vehicle-treated woozy mice, whereas GFAP immunoreactivity was markedly reduced following AAV treatment, approaching CT levels (Fig. 6). Thus, SIL1 gene therapy provided sustained protection against PC degeneration and associated cerebellar pathology.

**Figure 6.**
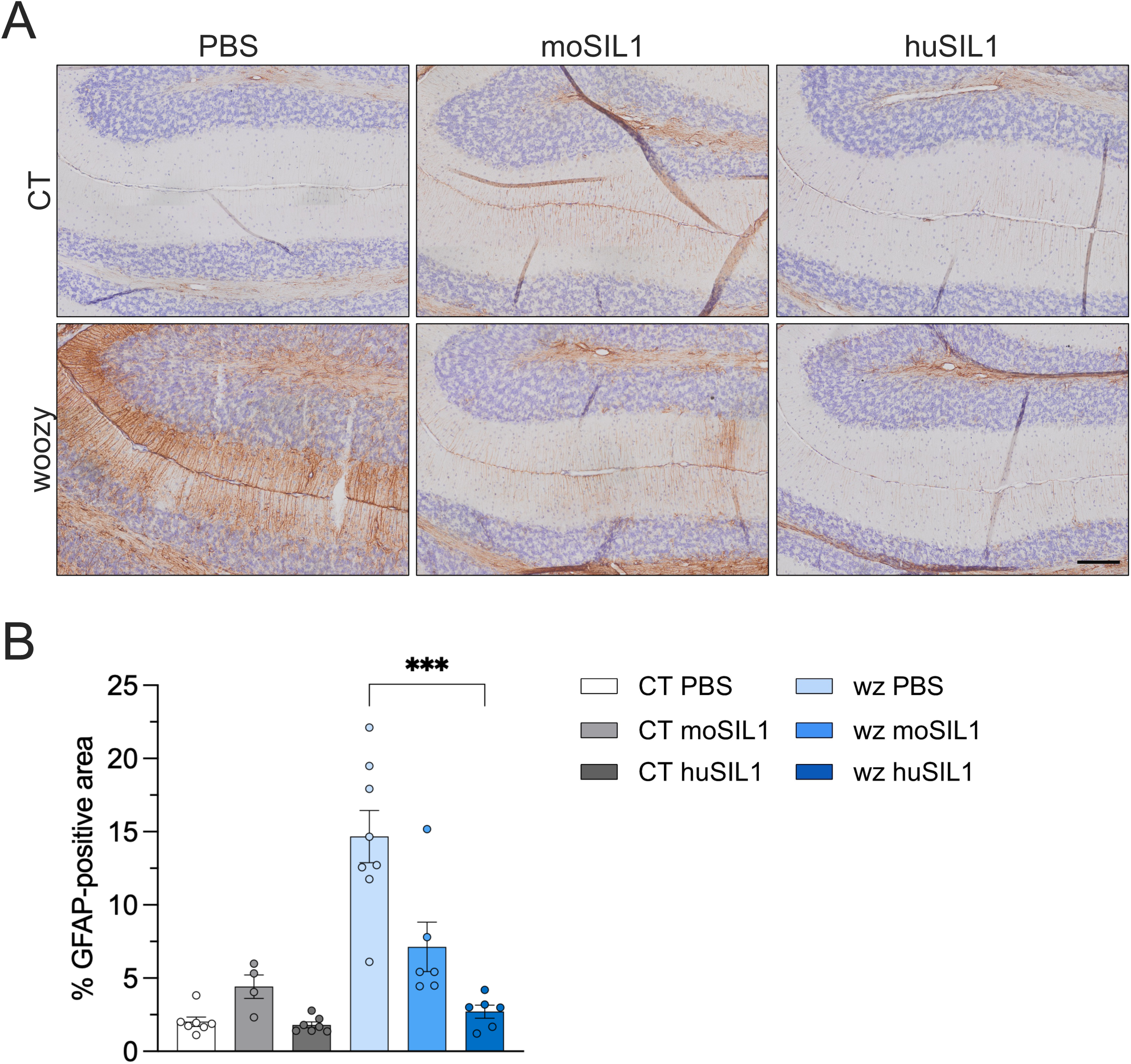
Intracerebroventricular injection of PHP.eB-CAG-moSIL1 and PHP.eB-CAG-huSIL1 reduces astrocytic activation. (**A**) Immunohistochemical analysis for GFAP in PBS, PHP.eB-CAG-moSIL1 or PHP.eB-CAG-huSIL1-treated CT or woozy mice at 26 weeks of age. Scale bar 100 μm. (**B**) Percentage of the GFAP positive area in the molecular layer of the cerebellar cortex. Data are the mean ± SEM of 3 brain sections from 4-8 mice per experimental group. *P* < 0.0001 by Kruskal-Wallis test; \*\*\**P* < 0.001 by Dunn’s post-hoc test.

### AAV treatment improves muscle strength and ameliorates ultrastructural muscle pathology

Woozy mice develop progressive skeletal muscle myopathy, first detectable by electron microscopy at 16 weeks and followed by impaired muscle function ^9,10^. Inverted-grid and grip-strength tests showed improved muscle resistance and strength in PHP.eB-CAG-SIL1-treated woozy mice compared with PBS-treated controls (Fig. 7), suggesting that icv AAV treatment also benefits skeletal muscle.

**Figure 7.**
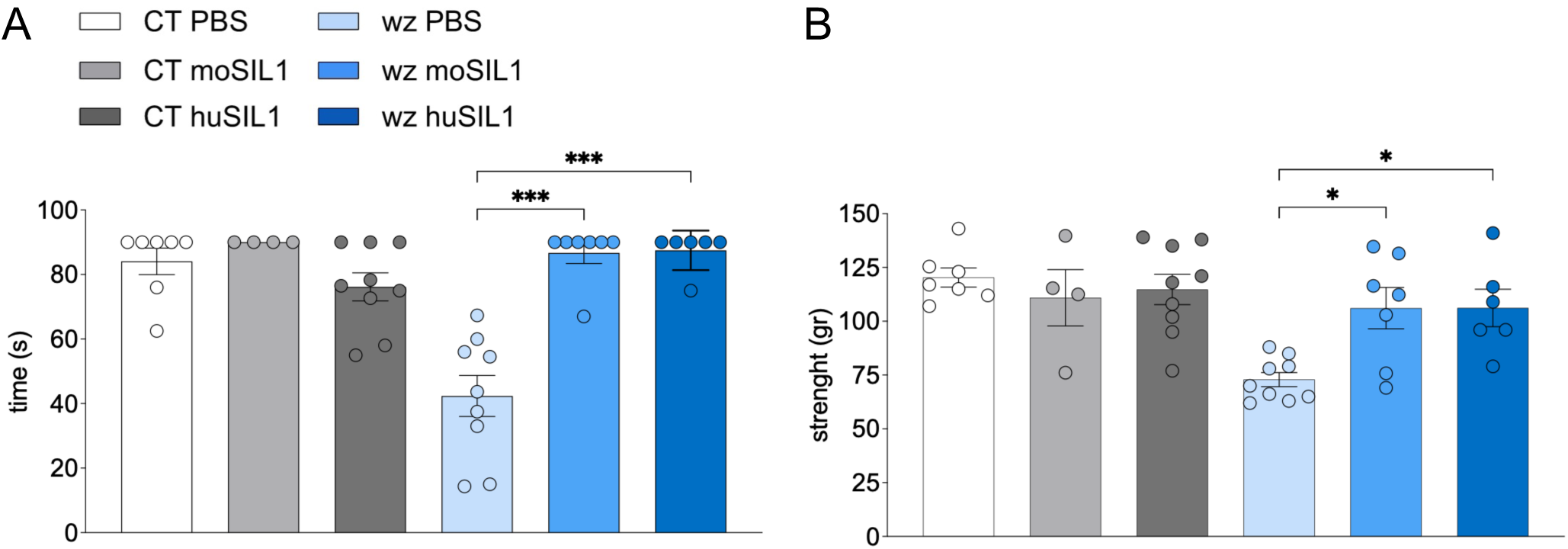
PHP-CAG-SIL1-injected woozy mice show improved muscle function. PBS, PHP.eB-CAG-moSIL1 and PHP.eB-CAG-huSIL1 treated animals were tested in the inverted grid (**A**) and grip strength (**B**) tests at 20 and 25 weeks of age, respectively. Data are the mean ± SEM. In (A), *P* < 0.0001 by Kruskal-Wallis test; \*\*\**P* < 0.001 by Dunn’s post hoc test. In (B), *P* = 0.0033 by Kruskal-Wallis test; \**P* < 0.05 by Dunn’s post-hoc test.

To assess this, we analysed the quadriceps muscles of the treated mice by histology and transmission electron microscopy (TEM). As reported previously ^10,18^, light microscopy examination found a high percentage of skeletal muscle fibres with non-subsarcolemmal nuclei in PBS-treated woozy mice, indicative of chronic myopathic remodelling associated with repeated cycles of degeneration and regeneration. This percentage was markedly reduced in woozy mice treated with AAVs encoding either mouse or human SIL1 (Fig. S11). Consistent with previous observations ^9,10^, TEM examination identified a number of ultrastructural alterations in the muscle fibres of PBS-treated woozy mice that were absent in controls. These included enlargements of the sarcoplasmic reticulum in the myofibrillar compartment and perinuclear autophagic vacuoles containing myelin-like membranous material (Fig. 8A, panels ii and vi). These alterations were significantly less marked in AAV-treated woozy mice (Fig. 8A, panels iii, iv, vii and viii), as documented by quantitative analysis (Fig. 8B).

**Figure 8.**
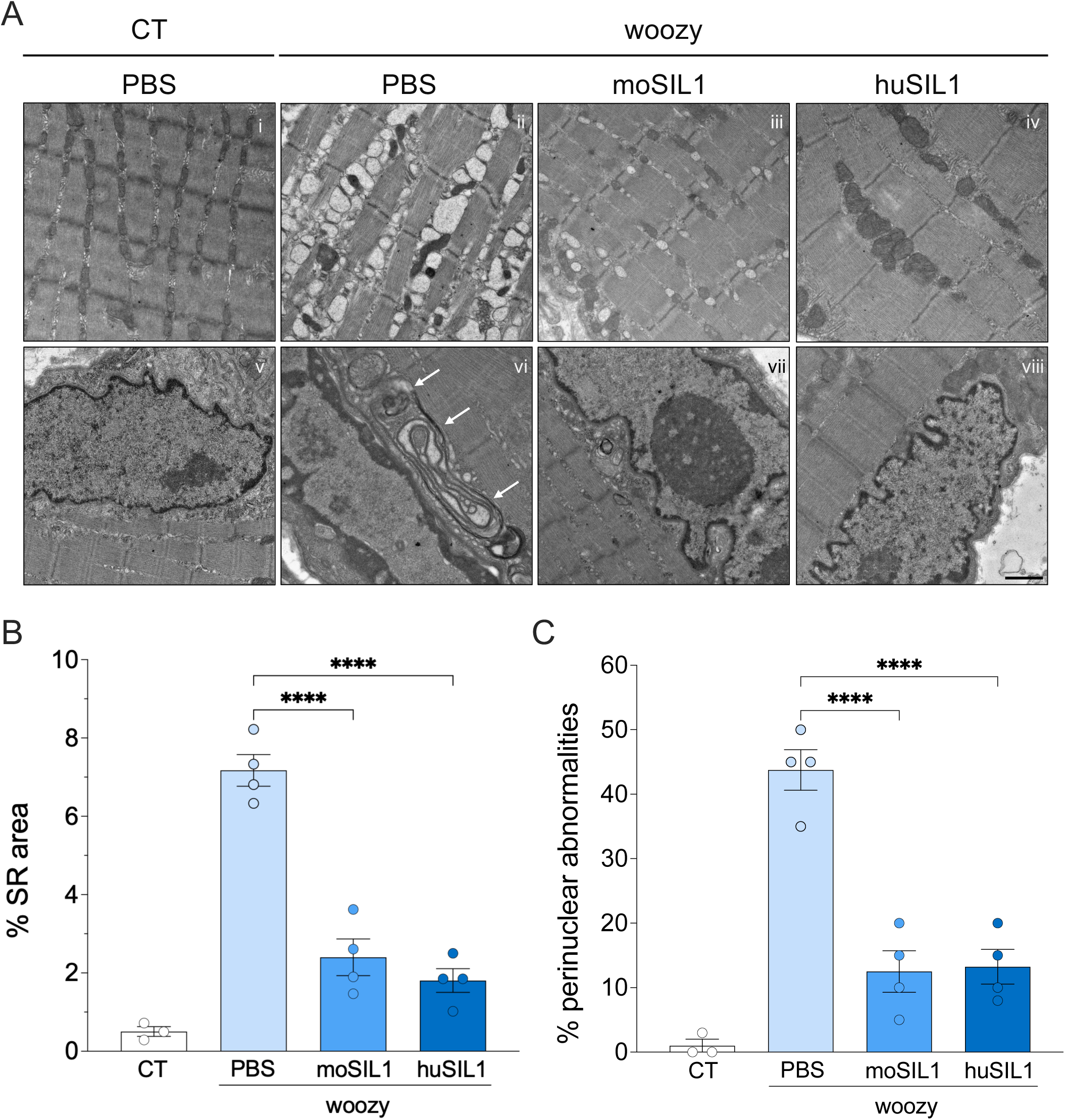
AAV-SIL1 administration attenuates the ultrastructural alterations in the quadriceps muscle fibres of woozy mice. (**A**) Quadriceps muscle ultrastructure of PBS- and AAV-treated mice at 26 weeks of age. Representative examples of normal myofibrillar (i) and perinuclear (v) muscle fibre ultrastructure in PBS-treated CT mice; pathological enlargement of the sarcoplasmic reticulum in the myofibrillar compartment of a woozy mouse treated with the vehicle (ii), or with AAVs encoding mouse (iii) or human (iv) SIL1; perinuclear autophagic vacuoles containing myelin-like membranous material (white arrows) in the skeletal muscle of a woozy mouse treated with PBS (vi) or AAVs encoding mouse (vii) or human (viii) SIL1. Scale bar, 1 μm. (**B**) Percentage of the myofibrillar area occupied by sarcoplasmic reticulum (SR)≤ in the different groups of mice. Data are mean ± SEM. F(3, 8) = 101.7, *P* < 0.0001 by two-way ANOVA; \*\*\*\**P* < 0.0001 by Dunnett’s post-hoc test. (**C**) Percentage of nuclei with perinuclear ultrastructural alterations. Data are mean ± SEM. F(3, 8) = 45.75, *P* < 0.0001 by two-way ANOVA; \*\*\*\**P* < 0.0001 by Dunnett’s post-hoc test.

Western blotting confirmed transgenic SIL1 expression in quadriceps muscle following icv PHP.eB-SIL1 administration (Fig. S12). Although lower than in the brain, SIL1 levels were comparable to endogenous levels in CT mice, consistent with previous reports that icv AAV administration can transduce peripheral tissues ^19,20^.

No evidence of treatment-related toxicity was observed. AAV-treated CT and woozy mice showed normal behaviour throughout the study, and serum chemistry parameters at 26 weeks were within the normal range (Table S1).

## DISCUSSION

The present study was undertaken to test a gene therapy approach for MSS, an ultrarare and highly debilitating disorder with onset in early infancy for which no disease-modifying therapy is currently available. We found that neonatal intracerebroventricular administration of AAV vectors encoding wild-type SIL1 prevents ataxia, attenuates cerebellar degeneration, as evidenced by preservation of Purkinje cells and reduced astrogliosis, and ameliorates skeletal muscle dysfunction and pathology in the woozy mouse model. Importantly, no evidence of toxicity associated with AAV administration or transgenic SIL1 overexpression was observed in either woozy or control mice. Selective restoration of SIL1 in Purkinje cells was sufficient to prevent cerebellar ataxia, suggesting a cell-autonomous contribution of Purkinje cell dysfunction to disease pathogenesis. These findings provide proof-of-principle evidence supporting AAV-mediated SIL1 gene therapy as a potential treatment strategy for MSS.

Currently, no effective treatment is available for MSS. Previous studies in woozy mice explored pharmacological approaches aimed at attenuating ER stress and maladaptive UPR signalling through modulation of the PERK pathway ^9,11,12^. Although these interventions produced partial benefit, they failed to completely rescue motor dysfunction or cerebellar degeneration, indicating that targeting downstream stress pathways may not be sufficient to counteract the consequences of SIL1 deficiency. A recent gene therapy approach used systemic AAV8 delivery resulting in predominantly liver-directed expression of a cell-penetrating SIL1-TAT fusion protein, with the liver serving as a “biofactory” for systemic production of the protein ^21^. SIL1-TAT was detected in skeletal muscle but not in the cerebellum, and treatment produced only modest functional benefits, delaying rather than preventing the onset of motor dysfunction, with only a limited delay of approximately 3 weeks ^21^. These observations strongly indicate that efficient delivery of SIL1 to the nervous system is required for robust therapeutic benefit. In this regard, the present study demonstrates that CNS-directed AAV-mediated gene therapy achieves substantially greater efficacy than previously explored pharmacological or peripheral gene delivery approaches.

A major finding of the present study is that AAV-mediated SIL1 delivery produced robust CNS expression and sustained rescue of ataxia-associated motor deficits through 26 weeks. Western blot analysis of cerebellar extracts showed SIL1 levels ranging from modestly above endogenous wild-type levels in moSIL1-treated mice to approximately sixfold higher in huSIL1-treated animals. However, because the antibody used for immunoblotting was raised against human SIL1, the stronger signal in huSIL1-treated mice may partly reflect higher antibody affinity for the human protein. ddPCR analysis of laser-capture microdissected PCs indicated approximately twofold SIL1 mRNA overexpression in the target neuronal population, consistent with robust but not excessively supraphysiological transgene expression.

Behavioural rescue was accompanied by substantial preservation of Purkinje cells, attenuation of molecular layer thinning, suppression of CHOP activation, and marked reduction of cerebellar astrogliosis, as evidenced by decreased GFAP immunoreactivity. These findings indicate that the functional benefit of AAV-SIL1 was associated with substantial preservation of cerebellar tissue and suppression of the pathological processes accompanying neurodegeneration. Although reactive astrocytosis is a well-established feature of neurodegenerative disorders, its contribution to MSS had not previously been explored. Because transgene expression was predominantly neuronal, the attenuation of astrogliosis is unlikely to result primarily from direct correction of astrocytes. Rather, it suggests that the glial response is secondary to neuronal degeneration and can be attenuated by protecting vulnerable Purkinje cells. Thus, SIL1 restoration appears to preserve not only neuronal survival but also aspects of cerebellar tissue homeostasis.

Preservation of Purkinje cells in AAV-treated woozy mice was substantial but not complete. Nevertheless, motor performance was essentially indistinguishable from that of heterozygous controls. This suggests that complete anatomical rescue may not be required to achieve robust functional benefit. Studies in several mouse models of cerebellar ataxia similarly show that partial preservation or functional restoration of PCs can substantially improve motor behaviour, likely reflecting the functional reserve and compensatory capacity of cerebellar circuits ^22–24^. Our findings therefore suggest that restoring SIL1 in a sufficient proportion of vulnerable PCs can preserve cerebellar network function and prevent overt ataxia despite residual neurodegeneration.

An additional important observation is the durability of the therapeutic effect. AAV-SIL1-treated woozy mice remained behaviourally indistinguishable from heterozygous controls throughout the 26-week observation period, the latest time point analysed. This sustained benefit is particularly relevant to MSS, in which neurological symptoms typically arise during infancy and evolve during childhood, although progression eventually stabilizes and life expectancy is usually not reduced. An effective therapy administered early in life could therefore provide long-lasting preservation of motor function and improve autonomy and quality of life. We also observed no overt adverse effects associated with neonatal icv AAV administration or sustained SIL1 expression in woozy or control mice, supporting the tolerability of AAV-mediated SIL1 transfer.

Another interesting finding of this study is that selective expression of SIL1 in PCs under the control of the L7-6 promoter was sufficient to prevent the development of ataxia. SIL1 deficiency likely renders PCs intrinsically vulnerable to chronic ER stress and defective protein homeostasis ^6,9^, and restoration of SIL1 expression specifically within these neurons is sufficient to markedly attenuate neurodegeneration and normalize cerebellar function in the woozy mouse model. These findings are consistent with observations in other inherited cerebellar ataxias, in which PCs represent a primary site of vulnerability and PC-directed genetic interventions substantially ameliorate motor deficits ^25–27^, underscoring the central contribution of PCs to cerebellar motor dysfunction. Our data therefore demonstrate that targeting PCs is sufficient to correct the cerebellar phenotype in the woozy mouse model and suggest that cell-specific gene therapy approaches may represent a viable strategy for some cerebellar disorders.

AAV-treated woozy mice also showed substantial improvements in muscle strength and endurance, as demonstrated by grip strength and inverted-grid tests. Consistent with these functional effects, histological and ultrastructural analyses showed attenuation of skeletal muscle pathology, including fewer myofibres with internal nuclei and reduced sarcoplasmic reticulum enlargement and perinuclear abnormalities. Transgenic SIL1 was detectable in skeletal muscle at levels similar to those in heterozygous mice, suggesting that even relatively modest restoration may provide significant benefit to muscle tissue.

Collectively, these findings support further development of AAV-mediated SIL1 gene therapy for MSS. Several questions, however, remain to be addressed. Although neonatal icv administration was highly effective in mice, clinical translation will require delivery strategies capable of efficiently transducing both CNS and peripheral tissues. Future studies should evaluate systemic and other administration routes using optimized AAV capsids and dosing regimens to achieve sufficient SIL1 expression in relevant target tissues while maintaining an acceptable safety profile.

Another important question for clinical translation is whether the therapeutic benefit observed with neonatal treatment can be extended to later stages of disease. Although prospective newborn genomic screening programs may eventually enable presymptomatic diagnosis and preventive treatment of severe genetic disorders ^28,29^, it will be important to determine whether AAV-SIL1 therapy remains effective after symptom onset, thereby defining the therapeutic window for intervention. Finally, although therapeutic benefit was maintained throughout the 26-week observation period and no overt toxicity was detected, longer-term studies and more comprehensive safety assessments will be required to evaluate the consequences of sustained SIL1 expression over the lifespan.

## METHODS

### Study design

This was a controlled, randomized laboratory study using the woozy (wz/wz) mouse model of MSS and heterozygous littermates as controls (CT). Based on previous findings showing no significant differences in motor performance or cerebellar anatomy between wild-type (wt/wt) and heterozygous (wt/wz) mice ^5,9,30^, heterozygous animals were used as controls throughout the study.

Newborn mice obtained from wt/wz × wz/wz matings (yielding approximately 50% CT and 50% woozy offspring) were randomly assigned at the litter level to receive AAV vectors or vehicle. Because sex cannot be reliably determined at birth, neonatal mice of both sexes were included and allocated to experimental groups irrespective of sex. Behavioural testing and histological and ultrastructural analyses were performed by investigators blinded to genotype and treatment. Short-term experiments were performed according to the experimental design shown in Fig. S5A to evaluate the effects of treatment on the onset of motor dysfunction and early PC degeneration. Mice were trained on the beam-walking test at 4 weeks of age and tested weekly until 9 weeks of age. The primary endpoint was performance in the beam-walking test, a sensitive measure of early impairments in motor coordination and balance in woozy mice ^9^. Sample size was determined by a priori power analysis. Based on previous data ^9^, assuming mean numbers of missteps of 2, 6, 16, 23, and 37 in vehicle-treated woozy mice at 5, 6, 7, 8, and 9 weeks of age, respectively, an AR(1) correlation of 0.50, and a residual variance of 2.00, six animals per group were required to detect a 60% reduction in missteps with a type I error of 0.05 and 90% power.

Long-term experiments were conducted according to the design shown in Fig. S5B. Animals were monitored until 26 weeks of age. In addition to beam walking, motor performance was evaluated using the accelerating rotarod, which detects motor impairment in woozy mice from approximately 9-10 weeks of age ^9^ and remains suitable for animals with advanced disease. Muscle function was assessed using the inverted grid and grip strength tests.

Animals were euthanized at the predefined experimental endpoints. At tissue collection, one hemisphere of each brain was immersion-fixed for histological, immunohistochemical, or ultrastructural analyses, whereas the contralateral hemisphere was snap-frozen for biochemical studies. Quadriceps muscles were processed similarly, with one muscle fixed for ultrastructural analyses and the contralateral muscle snap-frozen for histological and biochemical analysis ^9^.

### Ethic statement

Procedures involving animals and their care were conducted in conformity with the institutional guidelines at the Mario Negri Institute for Pharmacological Research IRCCS in compliance with national (D.lgs 26/2014; Authorization n. 19/2008-A issued March 6, 2008 by Ministry of Health) and international laws and policies (EEC Council Directive 2010/63/UE; the NIH Guide for the Care and Use of Laboratory Animals, 2011 edition). They were reviewed and approved by the Mario Negri Institute Animal Care and Use Committee that includes ad hoc members for ethical issues, and by the Italian Ministry of Health (Decreto no. D/07/2013-B and 301/2017-PR). Animal facilities meet international standards and are regularly checked by a certified veterinarian who is responsible for health monitoring, animal welfare supervision, experimental protocols, and review of procedures. All animal experiments were designed in accordance with the ARRIVE guidelines^31^, with a commitment to refinement, reduction and replacement, minimizing the numbers of mice, while using biostatistics to optimize mouse numbers.

### Generation of AAV vectors

Recombinant AAV vectors encoding mouse or human SIL1 under the control of the CAG promoter were generated by replacing the green fluorescent protein (GFP)-encoding sequence in the pAAV-CAG-GFP plasmid (Addgene #37825), with the cDNA encoding mouse or human SIL1. The mouse SIL1 coding sequence (GenBank: AJ297884.1), flanked by BamHI and HindIII restriction sites, was synthesized by GeneArt (Thermo Fisher Scientific). This construct included a synonymous A-to-C base substitution at position 264 from the ATG start codon to eliminate an internal BamHI restriction site. The human SIL1 coding sequence (GenBank: AJ297884.1), flanked by XbaI and HindIII restriction sites, was synthesized by GeneArt (Thermo Fisher Scientific) and cloned into the pAAV-CAG-GFP plasmid.

AAV vectors encoding moSIL1 or GFP under the control of the PC-specific promoter L7-6, an 844 bp fragment upstream of the start codon of exon 1B of the gene encoding the protein PCP2 (Purkinje specific protein 2, also known as L7) ^32^, were generated by replacing the CAG promoter with the L7-6 sequence synthesized by GeneArt and cloned into the NdeI and BamHI restriction sites. All vectors were sequenced (Microsynth) to verify the correctness of the sequence. AAV particles were generated by the AAV Vector Unit at ICGEB Trieste (www.icgeb.org/avu-core-facility) in HEK293T cells cultured in roller bottles by a three-plasmids transfection cross-packaging approach whereby the vector genome was packaged into AAV capsid. Viral particles were purified by PEG precipitation and two subsequent CsCl_2_ gradient centrifugations ^33^. The physical titer of recombinant AAVs was determined by absolute quantification of vector genomes (vg) packaged into viral particles, by real-time PCR using a standard curve of a plasmid containing the vector genome ^34^; values obtained were above 1x10^13^ vg/mL.

### Mice

Woozy mice (CXB5/By-Sil1wz/J) ^5^ were obtained from The Jackson Laboratory (Stock No. 003777). They were maintained by heterozygous mating and genotyped by standard PCR, as recommended by the supplier. They were housed at controlled temperature (22 ± 2°C) with a 12/12-hour light/dark cycle and free access to pelleted food and water. The health and home-cage behaviour of the treated mice were monitored daily, according to guidelines for health evaluation of experimental laboratory animals ^35^.

### Beam walking test

Mice were trained to walk along a metal beam 0.8 cm wide, 100 cm long, suspended 30 cm above bedding, for three days before testing. On the day of testing animals were given three trials. Mice were video-recorded and the number of hindfoot missteps and time to traverse the beam during the three trials were recorded by an investigator blinded to the experimental group. The average number of missteps and time to cross the beam during the three trials was used for statistical analysis.

### Accelerated Rotarod test

The accelerating Rotarod 7650 model (Ugo Basile) was used as described ^12^. Mice were trained three times before official testing. They were positioned on the rotating bar and allowed to become acquainted with the environment for 30 s. The rod motor was started at 7 rpm and accelerated to 40 rpm at a constant rate of 0.11 rpm/s for a maximum of 300 s. Performance was scored as latency to fall, in seconds. Animals were given three trials, and the average was used for statistical analysis.

### Tremor

Tremor was scored as mild (visible only when mice walked on the suspended beam) or severe (also visible when mice walked on a table top) by an investigator blinded to the experimental group.

### Inverted grid and grip strength tests

Muscle strength and endurance were assessed using the inverted grid and forelimb grip strength tests. For the inverted grid test, mice were placed on a wire grid that was then inverted and held approximately 40 cm above a padded surface. The latency to fall was recorded over a maximum duration of 90 s. Animals that fell before the end of the trial were tested up to three times, with 2-min intervals between trials. The mean latency across trials was used for statistical analysis.

Forelimb grip strength was measured using a grip strength meter consisting of a trapeze bar connected to a force transducer. Mice were allowed to grasp the bar with both forepaws and were then gently pulled backward by the tail until they released the bar. The peak force exerted was recorded in five consecutive trials separated by 30–60 s, and the mean value was used for statistical analysis.

### Intracerebroventricular AAV injection

AAV vectors were administered to newborn mice by icv injection a few hours after birth, and in all cases within 24 hours of life. Injection needles were prepared from flame-pulled glass capillaries and connected through polyethylene tubing (inner diameter 0.58 mm and 0.28 mm) to a Hamilton syringe. The injection apparatus was filled with the required volume of viral suspension (2 μL per ventricle). Pups were anesthetized by brief hypothermia on ice and positioned on a Petri dish placed over an LED light source, allowing visualization of the lateral ventricles through the skull. The needle was inserted approximately 1.5 mm deep at a point midway between the bregma and the eye, and both lateral ventricles were injected. Following the procedure, pups were placed in a clean cage in a cotton nest under a red heating lamp until recovery of normal body temperature and were then returned to the dam ^36^. The viral suspension contained 0.05% Evans Blue dye to facilitate visualization during injection.

### Immunofluorescence

Mouse embryonic fibroblasts grown on 13 mm-diameter glass coverslips were fixed in 4% paraformaldehyde (Sigma-Aldrich) for 10 min at room temperature (RT) and incubated in blocking/permeabilization solution containing 0.05% saponin, 0.5% bovine serum albumin (BSA), and 50 mM NH_4_Cl in PBS for 1h at RT. Cells were then incubated overnight at 4°C with primary antibodies (Table S2) diluted in blocking solution, washed three times in PBS, and incubated with Alexa Fluor 488- or 647-conjugated secondary antibodies for 1h at RT. Nuclei were stained with Hoechst 33258 (Invitrogen; 0.01 mg/mL in PBS) for 15 min at RT. Coverslips were mounted onto slides using ProLong Gold Antifade Reagent (Invitrogen), and cells were imaged using a Olympus FV-500 confocal microscope.

Cultured organotypic cerebellar slices were fixed in 4% paraformaldehyde (Sigma-Aldrich) for 4h at RT and then permeabilized free-floating in 0.5% Triton X-100 in PBS overnight at 4°C. Slices were blocked in 20% BSA in PBS for 4h at RT and subsequently incubated overnight at 4°C with primary antibodies diluted in 4% BSA in PBS. After three washes in PBS, slices were incubated with secondary antibodies diluted in 4% BSA in PBS for 2h at 4°C. Following three additional washes in PBS, nuclei were stained with Hoechst 33258 (Invitrogen; 0.01 mg/mL) in 4% BSA, 2% normal goat serum (NGS; Gibco), in PBS for 15 min at RT. Slices were mounted onto slides and coverslipped using ProLong Gold Antifade Reagent (Invitrogen), and imaged using an Olympus FV1000 confocal

Brains were sectioned on a cryostat (Leica CM1850) to obtain 30-μm parasagittal sections. Non-specific binding was blocked by incubating free-floating sections in blocking solution containing 10% horse serum (HS; Gibco), 1% Triton X-100 (Sigma), and 5% BSA (Sigma) for 2h at RT. Sections were then incubated overnight at 4°C with primary antibodies diluted in blocking solution. After three washes in PBS, sections were incubated with secondary antibodies diluted in blocking solution for 2h at RT. Nuclei were stained with Hoechst 33258 (Invitrogen; 0.01 mg/mL in PBS) for 10 min at RT. Sections were mounted onto slides and coverslipped using ProLong Gold Antifade Reagent (Invitrogen). Slides were imaged using an Evident Fluoview FV5000 confocal microscope.

### Immunohistochemistry

Sagittal brain sections (8 μm thick) were cut using a microtome (Leica RM2255). Tissue sections were mounted on positively-charged glass slides (Superfrost Plus, Thermo Fisher Scientific), incubated at 58°C for 20 min and then rehydrated by passing through an alcohol scale. An antigen retrieval step was done by washing samples in distilled water and boiling them in Antigen Decloaker Solution pH6 (Biocare Medical, diluted 1:10 in distilled H_2_O) for three 5-min cycles in a microwave at 750 W power with 1 min of cooling between each cycle. Samples were then equilibrated at room temperature, washed with distilled water, incubated in 3% H_2_O_2_ for 15 min to inhibit endogenous peroxidase activity, then incubated for 1h at room temperature in blocking solution (10% FBS (Sigma), 5% bovine serum albumin (Sigma), 1% Triton X-100 (Sigma) in PBS 1X (Gibco)). Sections were incubated at 4°C overnight with primary antibody (Table S2) diluted in blocking solution. Slides were then incubated with goat anti-rabbit Ig-G (H+L) Biotinylated secondary antibody (BA-1000, Vector Laboratories) 1:200 in blocking solution for 1h at room temperature, followed by 1h incubation with Avidin-Biotin Vectastain Elite ABC kit (Vector Laboratories) diluted 1:100 in PBS 1X. Finally, immunolabeling was visualized by application of the chromogen 3,3-diaminobenzidine (DAB) (Dako), followed by hematoxylin counterstaining (Mayer’s hematoxylin, Sigma-Aldrich).

Images were acquired with a VS120 Virtual Slide Microscope by Olympus and analyzed with NIH ImageJ software. Four sagittal sections were quantified for each mouse cerebellum using an algorithm to segment the stained area. The calbindin immunostained area in the cerebellar molecular layer was calculated as positive pixels/total assessed pixels. CHOP-positive cells were counted over the entire perimeter of the Purkinje cell layer. The width of the molecular layer of the cerebellar cortex was determined by averaging 15-20 independent measurements taken at points across lobules I-IX. ^37^. Slices were matched at equivalent midsagittal or parasagittal levels for comparisons among experimental groups.

### Histology

For skeletal muscle histological analysis, 10-μm-thick frozen sections of quadriceps femoris muscle were cut using a Leica CM 1850 UV cryostat and stained with haematoxylin (0.5%) and eosin G (1%). The percentage of non-subsarcolemmal muscle fibre nuclei was quantified from images acquired at 1000× magnification using a Zeiss Axioskop microscope. An average of 730 nuclei per muscle sample were analysed.

### Transmission electron microscopy

Quadriceps femoris muscle was excised with a razor blade, and fixed with 4% PFA and 2% glutaraldehyde in phosphate buffer 0.12 M, pH 7.4 for 4h at room temperature, followed by post-fixation with 1% O_s_O_4_ at room temperature in 0.12 M cacodylate buffer for 2h. After dehydration in graded series of ethanol, tissue samples were cleared in propylene oxide, embedded in Epoxy medium (Epon 812, Fluka) and polymerized at 60°C for 72h. From each sample, one semithin section (1 μm) was cut with a Leica EM UC6 ultramicrotome and mounted on glass slides for light microscopy. Ultrathin sections (60 nm thick) of areas of interest were obtained, counterstained with uranyl acetate and lead citrate, and examined with a Zeiss Libra 120 and a Hitachi HT7800 100kV transmission electron microscopes. The percentage of muscle fibre nuclei with perinuclear ultrastructural alterations was assessed by analysing 40 perinuclear areas in each animal. The percentage of the area occupied by the sarcoplasmic reticulum in the myofibrillar compartment was calculated by iTem software (Olympus Soft Imaging Solutions, Germany) in 20 random fields in each animal.

### Western blot

Proteins were separated according to their molecular weight by 10-12% SDS-PAGE and electro-transferred onto polyvinylidene fluoride or nitrocellulose membranes (Bio-Rad) by Trans-blot turbo (Bio-Rad). Protein transfer was checked by Ponceau S staining (Sigma-Aldrich) followed by washings in 100 mM Tris pH 7.5, 150 mM NaCl and 0.1% Tween 20 (TTBS) and blocking in 5% non-fat dry milk (NFDM, Sigma-Aldrich) in TTBS for 1h or in EveryBlot Blocking Buffer (Bio Rad 12010020) for 5 min at RT. Membranes were incubated overnight at 4°C or 1h at RT with the primary antibody (Table S2) diluted in 5% NFDM in TTBS, then rinsed three times with TTBS and incubated for 45 min with secondary antibody goat anti-mouse or anti-rabbit diluted conjugated with horseradish peroxidase (1:5,000; Bio-Rad). The chemiluminescent signal was revealed using Clarity Western ECL Substrate (Bio-Rad) and acquired by Chemidoc Imager MP (Bio-Rad). Quantitative densitometry of protein bands was done using ImageLab 6.0 software (Bio-Rad).

### Laser capture microdissection

Purkinje cells were isolated from the cerebellar cortex by laser capture microdissection (LCM). Cerebellar sections (20 μm thick) were cut using a cryostat (Leica CM1850) and collected on metal frame slides (Leica).

Slices were left in 75% cold ethanol for 2 min, then washed twice for 1 min in MilliQ water, incubated 1 min in cresyl violet dye (1% of powder in absolute ethanol, Merck Millipore), then held 5 sec in 90% ethanol, 5 sec in absolute ethanol, 1 min in fresh absolute ethanol, finally dried on absorbent paper and stored at -80°C with a dryer for later LCM. Before LCM, slices were equilibrated at -20°C, 4°C and room temperature for 15 min each. PCs were microdissected from the cerebellar cortex with a laser microdissector (Leica LMD7).

### Droplet digital PCR

Total RNA was extracted from LCM-isolated PCs using the RNeasy Mini Kit (Qiagen) and reverse-transcribed with the High-Capacity cDNA Reverse Transcription Kit (Applied Biosystems) using random primers. For each reaction, 0.3 ng of cDNA was added to 19 μL of QX200 EvaGreen Supermix (Bio-Rad) containing gene-specific primers at a final concentration of 100 nM (primer sequences and thermal cycling conditions are provided in Table S3). Droplets were generated using the QX200 Droplet Generator (Bio-Rad) according to the manufacturer’s instructions. PCR amplification was performed on a C1000 Touch Thermal Cycler (Bio-Rad). Droplets were analysed with a QX200 Droplet Reader (Bio-Rad), and absolute transcript abundance was quantified using QuantaSoft Analysis Pro software v1.0 (Bio-Rad). Data are presented as transcript copies per microliter of PCR reaction.

### Statistical analysis

Statistical analyses were done using Prism 11.0.2 (GraphPad Software). Normality of data distribution was assessed using the Shapiro–Wilk and Kolmogorov–Smirnov tests. Parametric data were analysed using two-way ANOVA or two-way repeated-measures (RM) ANOVA, as indicated in the figure legends, followed by Dunnett’s multiple-comparison test. Non-parametric data were analysed using the Kruskal–Wallis test followed by Dunn’s multiple-comparison test. Data are presented as mean ± SEM, with individual data points shown in all graphs. Differences were considered statistically significant at *P* < 0.05.

## Data availability

The data underlying this article are available in the article and in its online supplementary material.

## Acknowledgments

We are grateful to Maria Chiara Trolese and Davide Comolli for their assistance with icv injections and confocal microscopy. This work was funded by the European Union – NextGenerationEU, NRRP M6C2, Investment 2.1, *Enhancement and Strengthening of Biomedical Research in the NHS* (PNRR-MR1-2022-12375730), and by Fondazione Telethon and Fondazione Regionale per la Ricerca Biomedica (GMR25T2035). E.R. was the recipient of a Young Investigator Award (681848) from the National Ataxia Foundation.

## Author contributions

Conceptualization: RC

Investigation: CP, GL, AG, ER, AC, MS, LZ

Funding acquisition: RC, ER

Supervision: RC, FF

Writing – original draft: RC, CP, GL, AG,

Writing – review & editing: RC, CP, GL, AG, FF

