## Supplementary for "AAV-mediated SIL1 gene therapy prevents cerebellar degeneration and ataxia in a mouse model of Marinesco-Sjögren syndrome"

### Neonatal AAV-SIL1 gene therapy rescues Marinesco-Sjögren syndrome in mice

**The PDF file includes:**

Methods
Fig. S1 to S11
Table S1 to S3

**Other Supplementary Material for this manuscript includes the following:**

Movies S1 to S2
S1 Data. Excel spreadsheet containing, in separate sheets, the underlying numerical data for all Figure panels.

### Supplementary Methods

#### Mouse embryonic fibroblasts

Primary mouse embryonic fibroblasts were obtained from 13.5-14-day-old embryos as described *(39)*. Briefly, day E13.5 embryos from timed mating crosses were collected, and head, heart and liver, were removed and discarded. Tissue was thoroughly minced and incubated with 0.05% trypsin in PBS for 10 min at 37°C. Then 10 mg/mL DNAse I was added and incubated for other 10 min. Cells were dissociated by pipetting up and down 2-3 times and resuspended in Dulbecco's modified Eagle's medium (DMEM) supplemented with 10% FBS, 2 mM glutamine, 100 U/mL penicillin/streptomycin (Gibco). The cell suspension was passed through a 40-µm cell strainer and plated in a 100-mm tissue culture dish. Cells were incubated overnight; the medium was changed after 24h and incubated for another 1 or 2 days until confluence.

For AAV transduction, cells were plated (165.000 c/mL) on 24-well plates or 10-mm diameter glass coverslips coated with poly-L-lysine (Merck; 10 mg/mL overnight). After 24h cells were incubated with $10^{11}$-$10^{12}$ vg/mL of AAVs in OptiMEM (Gibco). After 24 h one volume of complete DMEM was added and cells were analyzed after additional 24h.

#### Cultured organotypic cerebellar slices

Cerebella were dissected from 9-10-day-old mice and transferred to a container with liquid agarose [2% low-melting point agarose in Geys Balanced Salt Solution with 33.33 mM glucose and 1 mM kynurenic acid (GBSSK)]. After cooling on ice, the agarose blocks were glued onto the vibratome disc, which was then placed in ice-cold GBSSK in the buffer reservoir of the vibratome (VT 1000S Leica). Several 350 µm sagittal slices were cut at low frequency (setting 6-7) and speed (setting 5-6). Slices were collected in a Petri dish containing ice-cold GBSSK and released from the agarose under a stereomicroscope using fine forceps. All subsequent steps were carried out under sterile conditions in a cell culture hood. Three to four slices were placed in a Millicell Cell Culture Insert (Millipore) in a six-well cell culture plate. Wells contained 1 ml of slice culture medium consisting of 50% MEM (Invitrogen), 22.5% HBSS, 25% horse serum (heat-inactivated; Gibco/Life Technologies), 50 U/ml penicillin, 50 µg/ml streptomycin (Invitrogen), 0.6% glucose, and 2 mM GlutaMax (Invitrogen). Slice cultures were kept under standard cell culture conditions (37°C, 5% $CO_2$/95% air) in a humidified atmosphere. The culture medium was changed every other day.

Slice cultures were exposed to $10^{12}$ vg/mL AAV 24 h after plating. At DIV 15 slices were fixed for immunofluorescence or lysed for WB analysis. Slices were examined with an Olympus Fluoview FV1000 confocal microscope.

#### Intracardiac perfusion and tissue processing

Mice injected with AAV vectors were perfused transcardially under deep anesthesia with 0.5× PBS (pH 7.4) at a flow rate of 10-12 mL/min for approximately 5-10 min. The brain was either frozen on dry ice and embedded in Optimal Cutting Temperature compound (OCT; Bio-Optica) for laser microdissection (LMD), or sectioned sagittally: one hemisphere was snap-frozen for biochemical analyses, whereas the other was processed for immunofluorescence or immunohistochemistry. For immunofluorescence, the hemisphere was immersion-fixed in 4% paraformaldehyde (Sigma, Cat. #158127) in PBS for approximately 48h, then washed in 0.5× PBS and transferred to 30% sucrose solution (D-sucrose, Carlo Erba, in PBS) for an additional 24-48 h. The tissue was subsequently frozen in n-pentane (Carlo Erba) at -45°C and stored at -80°C until use. For immunohistochemistry, the tissue was immersion-fixed in 10% formalin for at least 24h and then paraffin-embedded. Quadriceps muscle was snap-frozen in isopentane at -45°C and stored at -80°C.

**Preparation of tissue homogenates**

Brain homogenates were prepared at 10% (weight/volume) in PBS in the presence of protease inhibitors (Roche) and phosphatase inhibitors (PhosSTOP, Roche), using a glass-Teflon potter. The muscle tissue was homogenized using Turrax in RIPA buffer 1X [NaCl 75 mM, Triton X-100 0.5%, sodium deoxycholate (Sigma) 0.25%, SDS 0.05%, TrisHCl 25 mM] or in 1% SDS in $H_2O$ at 95°C, with protease and phosphatase inhibitors. Protein concentration was determined using the BCA assay (Pierce).

**Protein deglycosylation**

Cell or tissue extracts containing 20 µg of total protein were deglycosylated in a final volume of 20 µL. Samples were first denatured by adding one-tenth volume of 10× denaturing buffer (New England Biolabs) and incubating at 95°C for 10 min. Subsequently, one-tenth volume of 10% Nonidet P-40 and one-tenth volume of either 10× GlycoBuffer 2 (for PNGase F) or 10× GlycoBuffer 3 (for Endo H) (New England Biolabs), together with 500 U of either PNGase F or Endo H (500,000 U/mL; New England Biolabs), were added. Samples were then incubated at 37°C for 1 h or overnight. Reactions were stopped by adding 4× Laemmli sample buffer (Bio-Rad), followed by heating at 95°C for 10 min. Samples were then analyzed by western blotting.

### Supplementary Figures

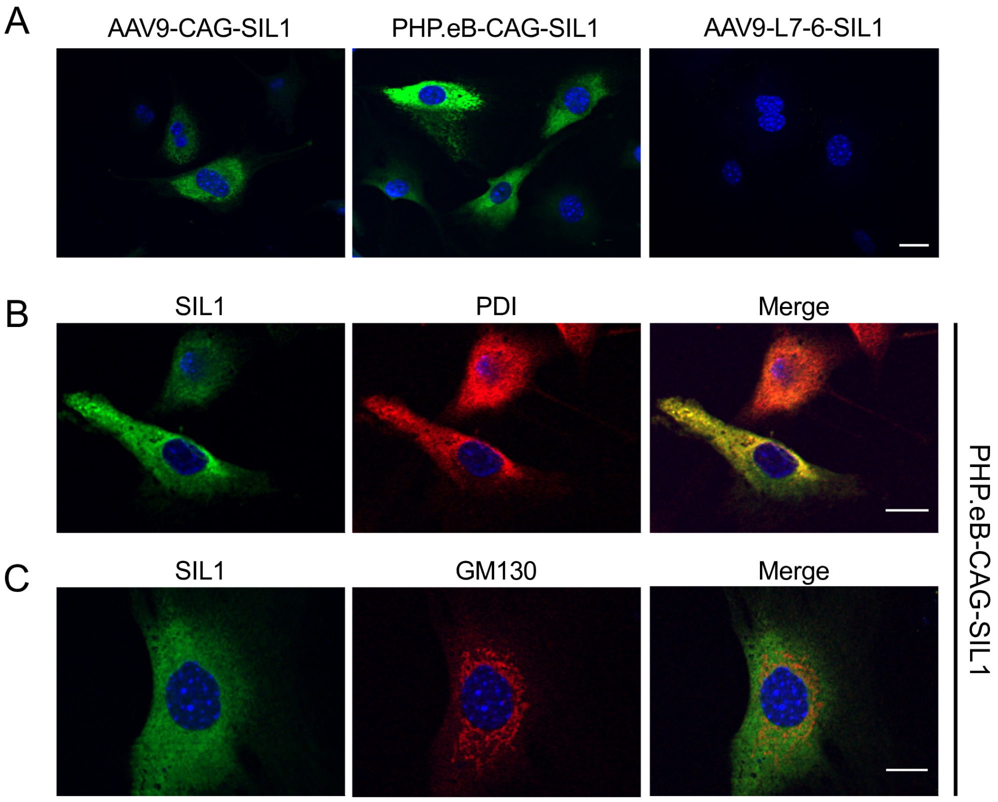


**Supplementary Fig. S1. SIL1 is specifically expressed in the ER of AAV-SIL1-tranduced woozy MEFs.** (**A**) Immunofluorescence analysis of woozy MEFs transduced with AAV9-CAG-moSIL1, PHP.eB-CAG-moSIL1 or PHP.eB-L7-6-moSIL1 with anti-SIL1 antibody (green). (**B**, **C**) Immunofluorescence of SIL1 (green) and PDI (B; red) or GM130 (C; red) in woozy MEFs transduced with PHP.eB-CAG-moSIL1. Cells were reacted with Hoechst 33258 to visualize the nuclei (blue). Scale bars: 20 µm in A, and 10 µm in B and C.

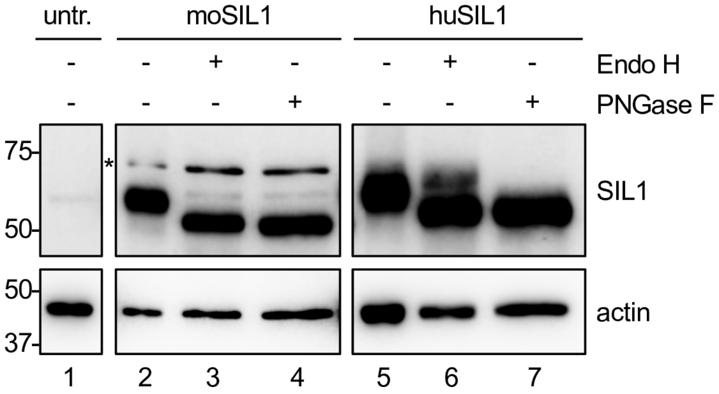


**Supplementary Fig. S2. AAV-encoded SIL1 is sensitive to Endo H and PNGase F deglycosylation.** MEF from woozy mice were left untreated or transduced with $10^{11}$ vg/mL PHP.eB-CAG-moSIL1 or PHP.eB-CAG-huSIL1- After 24h cells were lysed and the lysates were incubated without (-) or with (+) Endo H or PNGase F before Western blot analysis with anti-SIL1 and anti-actin antibodies. The asterisk indicates a non-specific band. Molecular weight markers are in kDa.

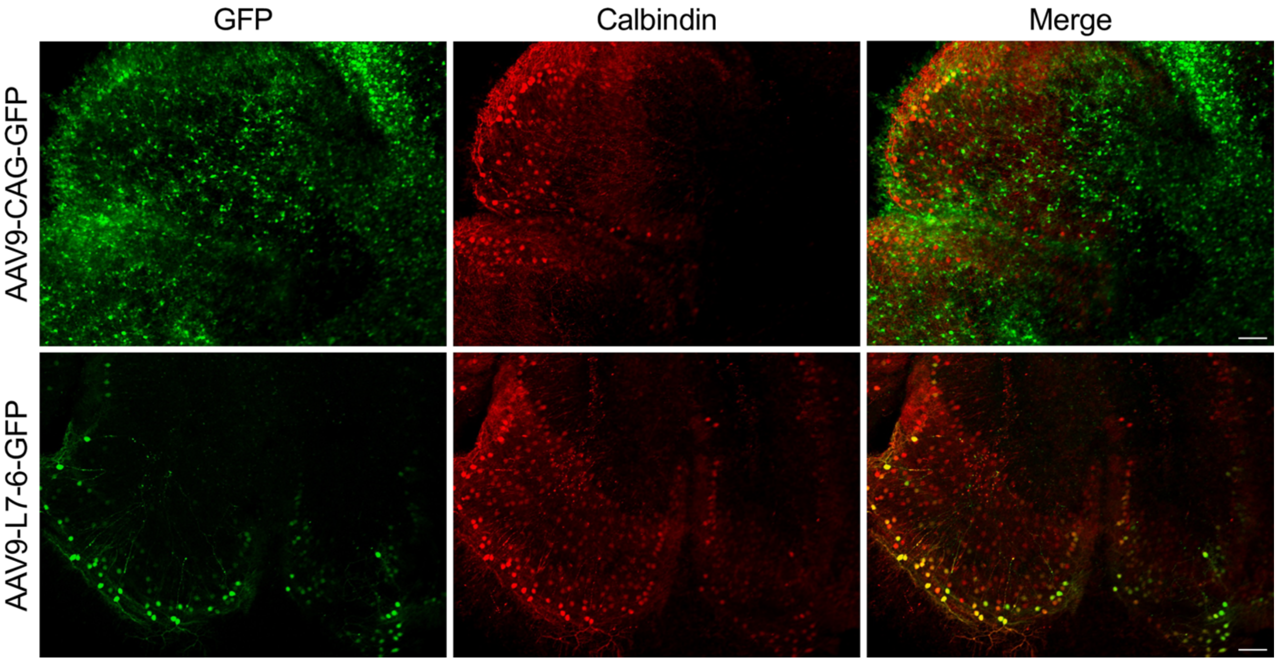


**Supplementary Fig. S3. Transduction of organotypic cerebellar slice cultures (COCS) with AAV9 vectors expressing GFP.** Cultured organotypic cerebellar slices transduced with AAV9-CAG-GFP or AAV9-L7-6-GFP were immunolabeled with an anti-calbindin antibody (red), which specifically labels Purkinje cells (PCs). GFP fluorescence is shown in green and calbindin immunofluorescence in red. Scale bars, 100 µm.

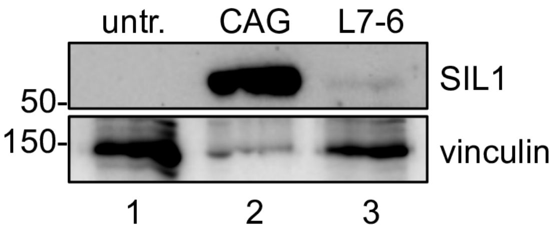


**Supplementary Fig. S4. PHP.eB viral vectors restore SIL1 expression in woozy COCS.** Western blot analysis of SIL1 and vinculin in woozy COCS left untreated (lane 1) or transduced with $1x10^{12}$ vg/mL PHP.eB-CAG-moSIL1 (lane 2) or PHP.eB-L7-6-moSIL1 (lane 3).

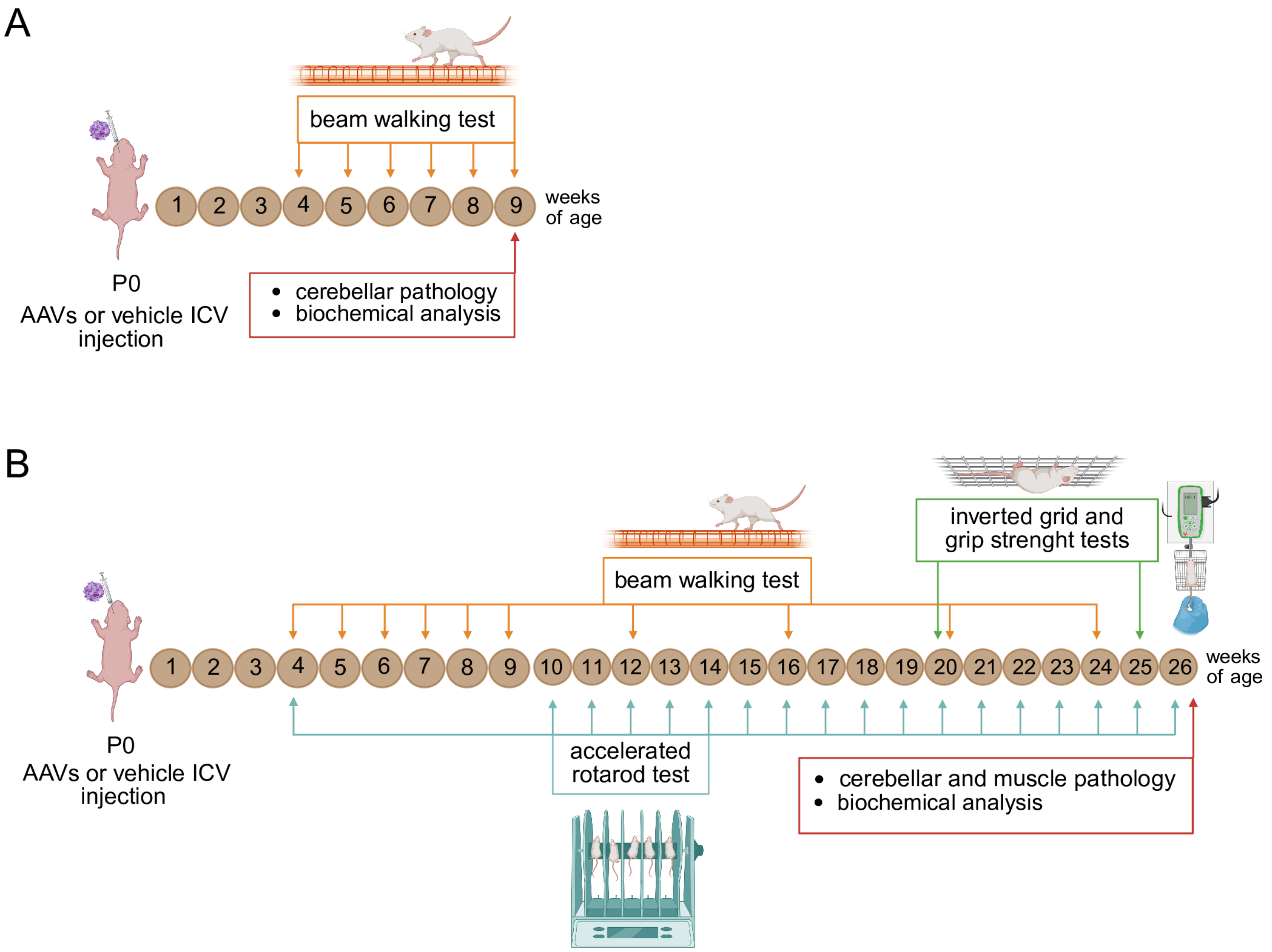


**Supplementary Fig. S5. Scheme of mouse treatment and analysis.** (**A**) Short-term experiment. (**B**) Long-term experiment.

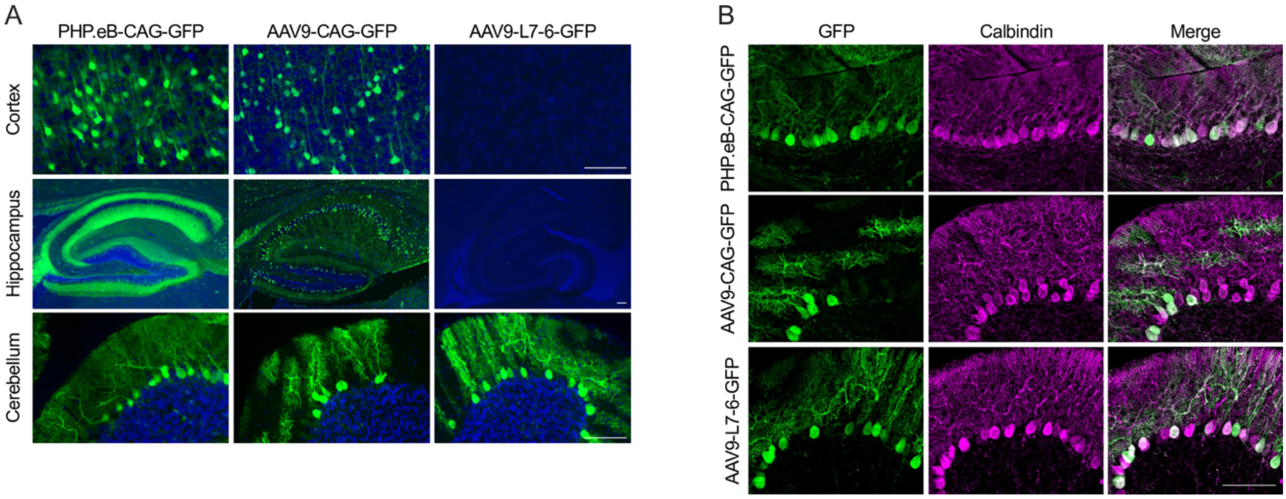


**Supplementary Fig. S6. Analysis of GFP expression in mice following intracerebroventricular AAV injection.** (**A**) Immunostaining for GFP in representative brain regions of mice injected icv with the indicated AAV vectors and analyzed 4 weeks post-injection. Scale bars, 100 µm. (**B**) Immunostaining for GFP and calbindin in the cerebellum of the same mice shown in (A). Scale bar, applicable to all panels, 50 µm.

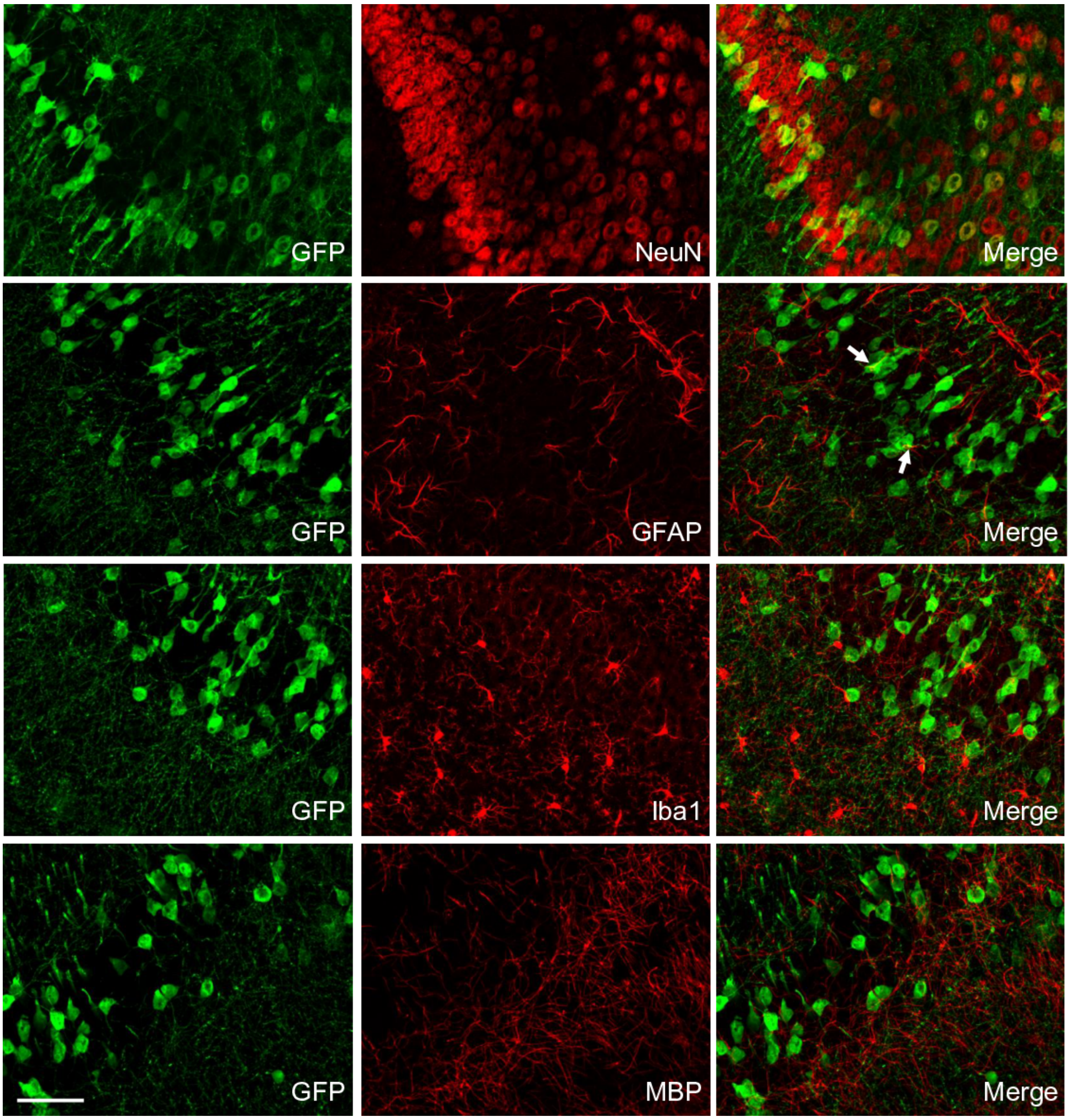


**Supplementary Fig. S7. Intracerebroventricular AAV injection results in predominantly neuronal transgene expression.** Immunostaining for GFP together with NeuN (neuronal marker), GFAP (astrocytic marker), Iba1 (microglial marker), or MBP (oligodendrocyte marker) in the hippocampus of mice injected with AAV9-CAG-GFP and analyzed 4 weeks post-injection. Arrows indicate GFP-positive astrocytes. Scale bar, applicable to all panels, 50 μm.

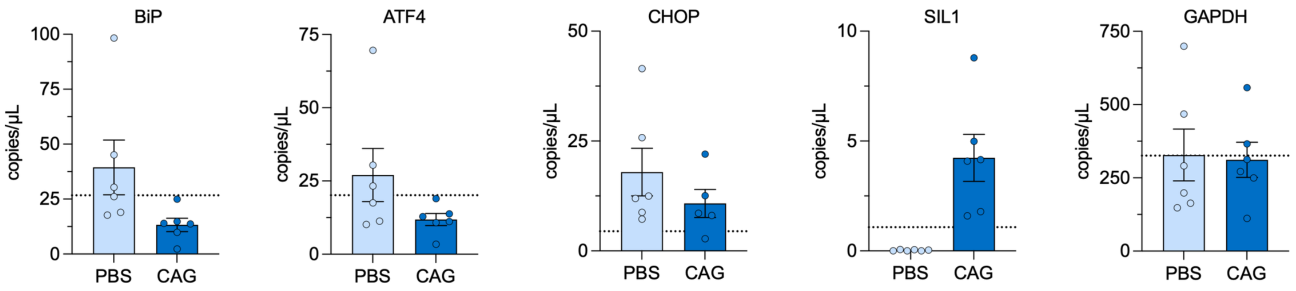


**Supplementary Fig. S8. Expression of UPR genes is reduced in Purkinje cells of PHP.eB-CAG-moSIL1-treated woozy mice.** Quantification of mRNA levels of the indicated genes by ddPCRs in LCM-isolated Purkinje cells from PBS- and PHP.eB-CAG-moSIL1-treated woozy mice at 6 weeks of age. Data are shown as mean ± SEM. Horizontal dotted lines indicate the mean levels measured in heterozygous PBS-treated mice (n = 5).

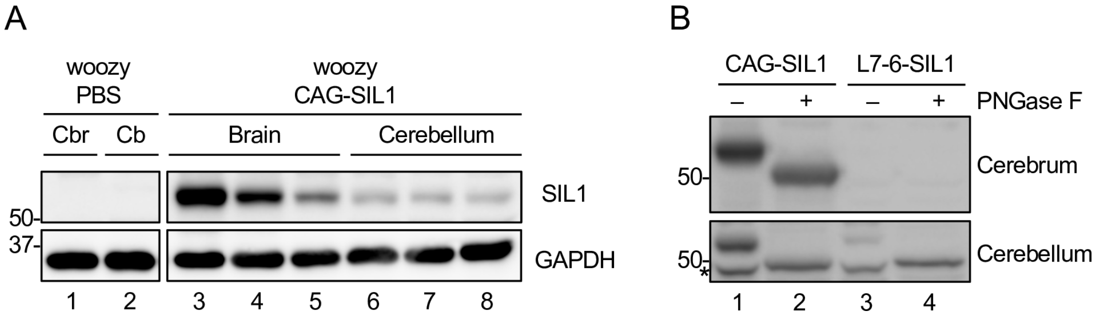


**Supplementary Fig. S9. Western blot analysis of SIL1 in AAV-treated woozy mice.** (**A**) Western blot analysis of SIL1 and GAPDH in the cerebrum (Cbr) and cerebellum (Cb) of PBS- and PHP.eB-CAG-moSIL1 (CAG-SIL1)-treated woozy mice. (**B**) Western blot analysis of SIL1 in the cerebrum and cerebellum of woozy mice treated with PHP.eB-CAG-moSIL1 (CAG-SIL1) or PHP.eB-L7-6-moSIL1 (L7-6-SIL1), before and after PNGase F treatment. Molecular weight markers are shown in kDa. Note that the deglycosylated SIL1 band co-migrates with a non-specific band in cerebellar extracts (indicated by the asterisk).

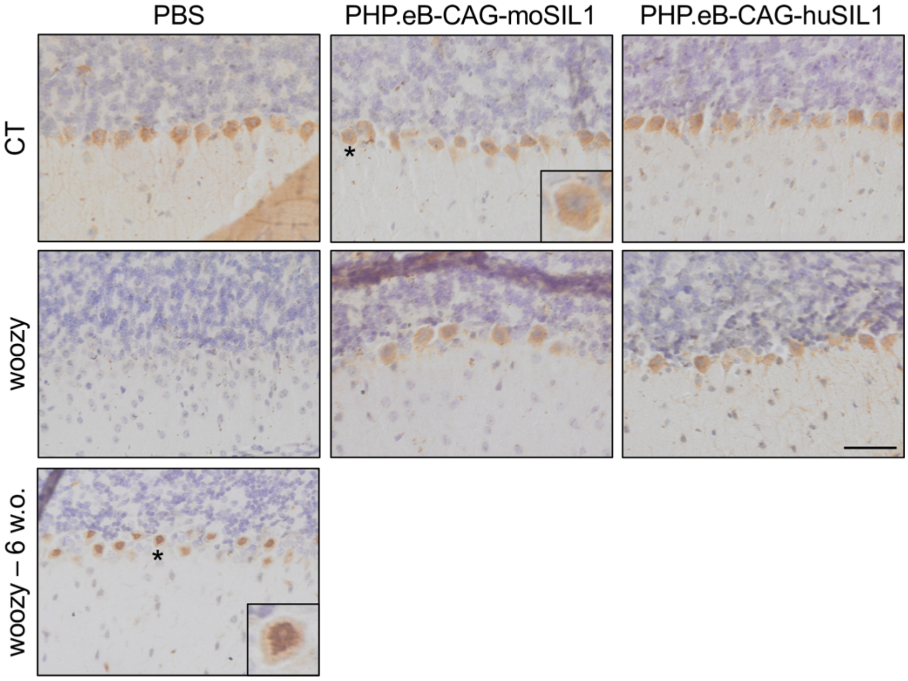


**Supplementary Fig. S10. AAV-SIL1-treated mice show no CHOP positivity in rescued Purkinje cells.** Immunohistochemical analysis of CHOP in CT and woozy mice treated with PBS, PHP.eB-CAG-moSIL1, or PHP.eB-CAG-huSIL1 and analyzed at 26 weeks of age. An untreated 6-week-old woozy mouse was stained in parallel as a positive control. Asterisks indicate the cells magnified in the inset, illustrating the difference between cytoplasmic background staining in a CHOP-negative cell and the characteristic nuclear staining of a CHOP-positive Purkinje cell. Scale bar, 50 µm.

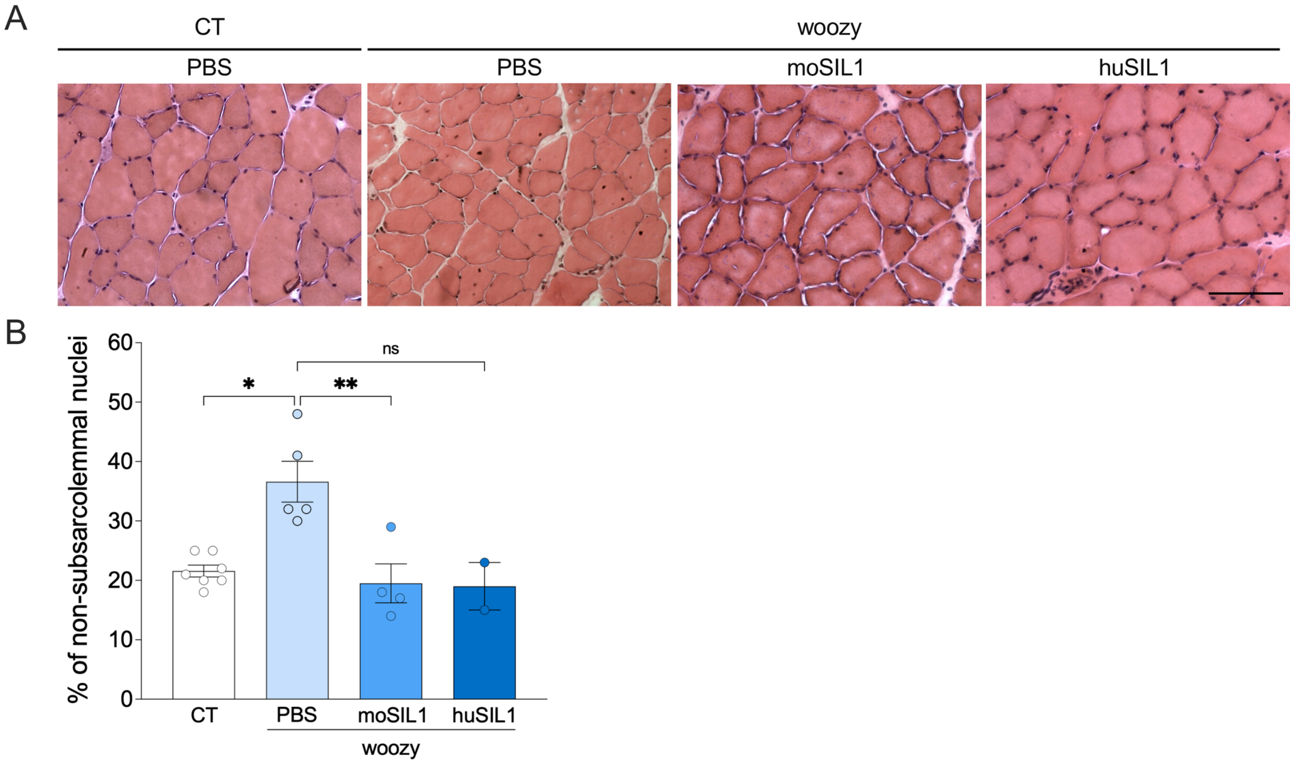


**Supplementary Fig. S11. AAV-SIL1 administration reduces the percentage of non-subsarcolemmal nuclei in the quadriceps muscle fibers of woozy mice.** (**A**) Hematoxylin and eosin staining of quadriceps muscle of PBS- and AAV-treated mice at 26 weeks of age. Scale bar, 100 μm. (**B**) Percentage of non-subsarcolemmal muscle fiber nuclei. Data are mean ± SEM. P = 0.0019 by Kruskal-Wallis test; **P* < 0.05, ***P* < 0.01, ns, not significant (P = 0.0746) by Dunn's post-hoc test.

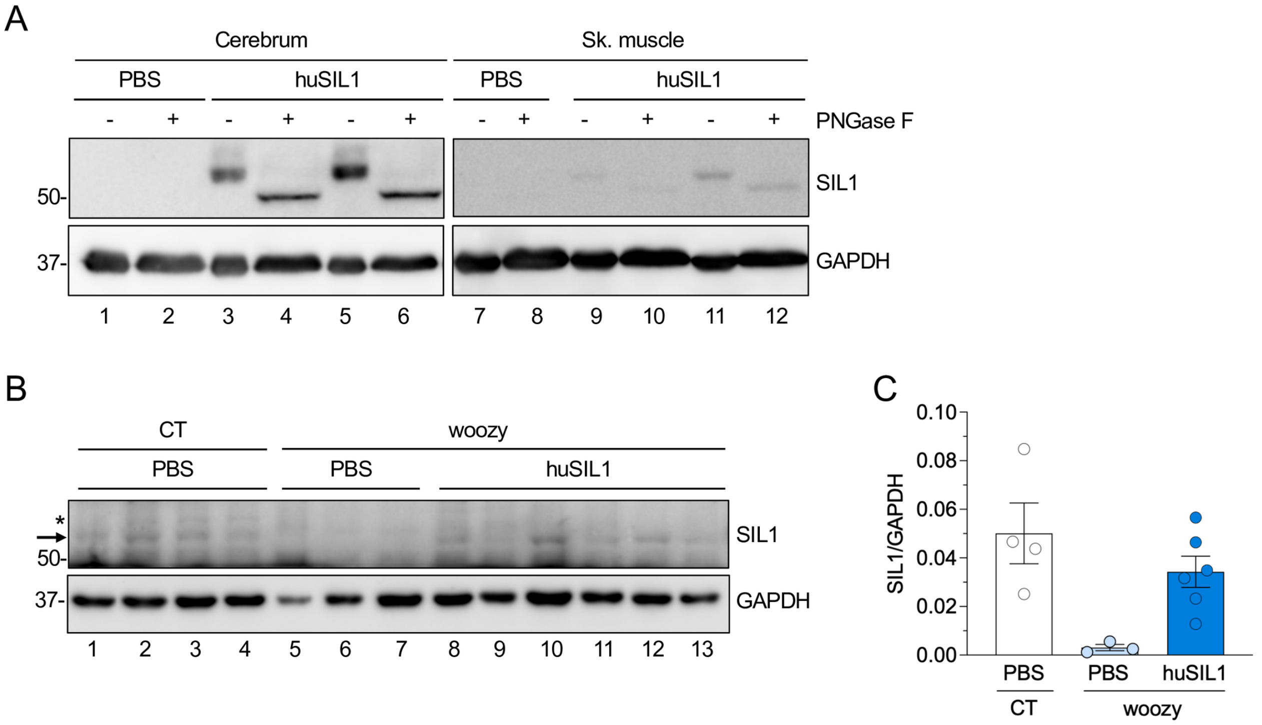


**Supplementary Fig. S12. Analysis of transgenic SIL1 expression in the brain and skeletal muscle of AAV-treated woozy mice.** (**A**) Western blot analysis of SIL1 and GAPDH in the indicated tissues from woozy mice injected icv at P1 with PBS or PHP.eB-CAG-huSIL1 and analyzed at 26 weeks of age, before (-) and after (+) deglycosylation with PNGase F. (**B**) Western blot analysis of SIL1 in the quadriceps muscles from CT mice injected with vehicle (PBS) and woozy mice injected with PBS or PHP.eB-CAG-huSIL1. The asterisk indicates a non-specific band, and the arrow indicates the specific SIL1 signal (**C**) SIL1 levels were quantified by densitometric analysis of the blot shown in (B) and normalized to GAPDH levels. Data are presented as mean ± SEM.

### Supplementary Tables

**Table S1. Serum chemistry parameters in control (CT) and woozy mice at 26 weeks of age.** Serum chemistry parameters measured using the Element RC General Health Panel (Scilvet). Data are the mean ± SEM for 2–3 animals per experimental group, except for the CT PBS group, in which only one animal was analyzed. ALB, albumin; TP, total protein; GLOB, globulin; A/G, albumin-to-globulin ratio; TBIL, total bilirubin; GGT, gamma-glutamyl transferase; AST, aspartate aminotransferase; ALT, alanine aminotransferase; ALP, alkaline phosphatase; CREA, creatinine; BUN, blood urea nitrogen; BUN/CREA, blood urea nitrogen-to-creatinine ratio; GLU, glucose; TC, total cholesterol; Ca, calcium; PHOS, inorganic phosphorus; K, potassium; Na, sodium; n.d., not detected.

| | **Reference values** | **CT PBS** | **CT huSIL1** | **CT moSIL1** | **woozy PBS** | **woozy moSIL1** |
|---|---|---|---|---|---|---|
| **ALB** | 3-4 g/dL | 3.1 | 3.2±0.07 | 3.13±0.06 | 3.1±0.17 | 2.33±1.15 |
| **TP** | 5-7 g/dL | 4.7 | 5.45±0.07 | 5.17±0.21 | 5.067±0.35 | 3.97±2.16 |
| **GLOB** | 1.5–2.5 g/dL | 1.6 | 2.15±0.07 | 2.033±0.15 | 1.93±0.21 | 1.63±0.93 |
| **A/G** | - | 1.87 | 1.52 ± 0.08 | 1.54±0.12 | 1.62±0.14 | 1.5±0.21 |
| **TBIL** | 0.1-0.9 mg/dL | 0.1 | n.d. | 0.133±0.06 | 0.1±0 | 0.13±0.06 |
| **GGT** | 0-10 U/L | < 2 | < 2 | < 2 | < 2 | < 2 |
| **AST** | 77-383 U/L | 81 | 57±11.31 | 66.33±12.2 | 70.33±16.92 | 80±22.63 |
| **ALT** | 40-189 U/L | 52 | 47±11.31 | 41.66±9.07 | 43.33±2.52 | 42.33±26.41 |
| **ALP** | 66-262 U/L | 47 | 46±9.9 | 44.33±3.22 | 41±4 | 34±14.73 |
| **CREA** | 0.0-0.5 mg/dL | 0.1 | 0.2±0.14 | 0.4±0.14 | 0.33±0.12 | 0.2±0.0 |
| **BUN** | 21-26 mg/dL | 19.19 | 20.82±2.16 | 19.26±1.67 | 21.25±2.61 | 15.72±8.92 |
| **BUN/CREA** | - | 191.9 | 143.92±112.55 | 50.33±11.89 | 69.39±25.05 | 78.6±44.60 |
| **GLU** | 196-278 mg/dL | 112.13 | 134.2±2.21 | 112.41±11.94 | 116.05±15.49 | 109.61±68.99 |
| **TC** | 70-140 mg/dL | 106.08 | 150.59±29.42 | 126.03±39.03 | 124.13±20.98 | 94.8±45.03 |
| **Ca** | 7.9-10.5 mg/dL | 9.19 | 9.33±0.04 | 9.37±0.11 | 9.05±0.35 | 7.13±4.11 |
| **PHOS** | 5.6-9.2 mg/dL | 7.12 | 7.13±0.98 | 7.34±0.56 | 7.00±0.47 | 5.62±3.10 |
| **K** | 5.3-6.3 mmol/L | 6.64 | 7.14±1.06 | 6.9±0.38 | >8 | 7.54±0.4 |
| **Na** | 138-186 mmol/L | 148.9 | 148.55±0.5 | 148.87±0.48 | 148.7±0.4 | 144.73±7.22 |

**Table S2. List of the antibodies used for Western blot (WB), Immunofluorescence (IF) and immunohistochemistry (IHC).**

| Antigen | Clone | Synonyms | Company | Code | Species | Dilution (application) |
|---|---|---|---|---|---|---|
| Calbindin | D28-K | | Swant | CB38 | Rabbit | 1:1000 (IF)<br>1:2500 (IHC) |
| Calbindin | D28-K | | Sigma | C9848 | Mouse | 1:1000 (WB) |
| CHOP | | DNA damage-inducible transcript 3 DDIT3 | Santa Cruz | sc7351 | Mouse | 1:250 (IHC) |
| GAPDH | | Glyceraldehyde-3-phosphate dehydrogenase | Sigma | G8795 | Mouse | 1:40000 (WB) |
| GFAP | | Glial fibrillary acidic protein | Merck-Millipore | MAB3402 | Mouse | 1:100 (IF) |
| GFAP | | Glial fibrillary acidic protein | Dako | Z0334 | Rabbit | 1:2500 (IHC) |
| GFP | | Green fluorescent protein | Novus | NB600-308 | Rabbit | 1:1000 (WB, IF) |
| GFP | | Green fluorescent protein | Genetex | GTX13970 | Chicken | 1:1000 (IF) |
| GM130 | | | BD Transduction | 610823 | Mouse | 1:500 (IF) |
| Iba1 | | Ionized calcium-binding adapter molecule 1 | Wako | 019-19741 | Rabbit | 1:100 (IF)<br>1:1000 (IHC) |
| MBP | D8X4Q | Myelin basic protein | Cell signaling | 78896 | Rabbit | 1:500 (IF) |
| NeuN | | | Merck-Millipore | mab377 | Mouse | 1:100 (IF) |
| PDI | RL90 | P4HB; Protein disulfide-isomerase | Abcam | Ab2792 | Mouse | 1:100 (IF) |

**Table S3. Primer sequences and thermal protocol for ddPCR.**

| Primer name | Sequence | Thermal protocol |
|---|---|---|
| ATF4 Fw | 5'-GAGCTTCCTGAACAGCGAAGTG-3' | 1) 95°C – 5 min<br>2) 95°C – 30 s<br>3) 64°C – 1 min<br>4) GO TO step 2 x 50<br>5) 4°C – 5 min<br>6) 90°C – 5 min<br>7) 4°C infinite |
| ATF4 Rv | 5'-TGGCCACCTCCAGATAGTCATC-3' | |
| CHOP Fw | 5'-CCACCACACCTGAAAGCAGAA-3' | |
| CHOP Rv | 5'-AGGTGCCCCCAATTTCATCT-3' | |
| BIP Fw | 5'-ACTCCGGCGTGAGGTAGAAA-3' | |
| BIP Rv | 5'-AGAGCGGAACAGGTCCATGT-3' | |
| GAPDH Fw | 5'-TGTTTGTGATGGGTGTGAACC-3' | |
| GAPDH Rv | 5'-CTGTGGTCATGAGCCCTTCC-3' | |
| Sil1_Exon7 Fw | 5'-CCTGGTGAAGGAGTATGCTGCTTT-3' | 1) 95°C – 5 min<br>2) 95°C – 30 s<br>3) 58°C – 1 min<br>4) GO TO step 2 x 50<br>5) 4°C – 5 min<br>6) 90°C – 5 min<br>7) 4°C infinite |
| Sil1_Exon10 Rv | 5'-TTCAGGAACTGCTGCTGGGCATAG-3' | |
